# Predictors of human-infective RNA virus discovery in the United States, China and Africa, an ecological study

**DOI:** 10.1101/2021.09.13.460031

**Authors:** Feifei Zhang, Margo Chase-Topping, Chuan-Guo Guo, Mark E.J. Woolhouse

## Abstract

**Background:** The variation in the pathogen type as well as the spatial heterogeneity of predictors make the generality of any associations with pathogen discovery debatable. Our previous work confirmed that the association of a group of predictors differed across different types of RNA viruses, yet there have been no previous comparisons of the specific predictors for RNA virus discovery in different regions. The aim of the current study was to close the gap by investigating whether predictors of discovery rates within three regions—the United States, China and Africa—differ from one another and from those at the global level.

**Methods:** Based on a comprehensive list of human-infective RNA viruses, we collated published data on first discovery of each species in each region. We used a Poisson boosted regression tree (BRT) model to examine the relationship between virus discovery and 33 predictors representing climate, socio-economics, land use, and biodiversity across each region separately. The discovery probability in three regions in 2010–2019 was mapped using the fitted models and historical predictors.

**Results:** The numbers of human-infective virus species discovered in the United States, China and Africa up to 2019 were 95, 80 and 107 respectively, with China lagging behind the other two regions. In each region, discoveries were clustered in hotspots. BRT modelling suggested that in all three regions RNA virus discovery was best predicted by land use and socio- economic variables, followed by climatic variables and biodiversity, though the relative importance of these predictors varied by region. Map of virus discovery probability in 2010– 2019 indicated several new hotspots outside historical high-risk areas. Most new virus species since 2010 in each region (6/6 in the United States, 19/19 in China, 12/19 in Africa) were discovered in high risk areas as predicted by our model.

**Conclusions:** The drivers of spatiotemporal variation in virus discovery rates vary in different regions of the world. Within regions virus discovery is driven mainly by land-use and socio- economic variables; climate and biodiversity variables are consistently less important predictors than at a global scale. Potential new discovery hotspots in 2010–2019 are identified. Results from the study could guide active surveillance for new human-infective viruses in local high risk areas.

**Funding:** Darwin Trust of Edinburgh; European Union.

## Introduction

RNA viruses are the primary cause for emerging infectious diseases with epidemic potential, given that they have high rate of evolution and high capacity to adapt to new hosts (Woolhouse et al., 2016). In recent decades, infectious diseases caused by severe acute respiratory syndrome coronavirus (SARS-CoV), Middle East respiratory syndrome coronavirus (MERS-CoV), Bundibugyo Ebola virus and SARS-CoV-2 present major threats to the health and welfare of humans (Albarino et al., 2013; Ksiazek et al., 2003; Mackay et al., 2015; World Health Organisation, 2020). Detection of formerly unknown human-infective RNA viruses in the earliest stage after the emergence are essential for controlling the infections they cause. Measures to implement early detection include not only advanced diagnostic techniques (Lipkin et al., 2013), but more importantly the idea where to look for them (so-called hotspots) (Morse et al., 2012).

Socio-economic, environmental, and ecological factors related to both virus natural history and research effort have been found to affect the discovery of emerging RNA viruses (Jones et al., 2008; Morse et al., 2012; Rosenberg, 2015; Zhang et al., 2020). However, these factors are highly spatially heterogeneous, making the generality of any associations with discovery debatable. For example, the United States, China, and Africa have experienced different rates of socio-economic, environmental, and ecological changes in the last one hundred years. The United States has always had better resources to discover new viruses. For example, the Rockefeller Foundation—a U.S. foundation—supported the discovery of 23 arboviruses in Latin America, Africa, and India in 1951–1969 (Rosenberg, 2015). China has seen urban land coverage more than double and GDP per capita increase by seven times since the 1980s (Ritchie, 2018; Roser, 2013). Nine out of 223 human-infective RNA viruses have been originally discovered in China, and all were discovered after 1982 (Zhang et al., 2020). In contrast, effective surveillance is challenging in less developed regions such as large parts of Africa given resource constraints (Petti et al., 2006).

There have been no previous comparisons of the specific predictors for RNA virus discovery in different regions. In this study, we applied a similar methodology from our previous study of global patterns of discovery of human-infective RNA viruses (Zhang et al., 2020) to investigate whether predictors of discovery rates within three regions—the United States, China and Africa—differ from one another and from those at the global level, using three new virus discovery data sets. We also mapped discovery probability in three regions in 2010–2019 using the fitted models and historical predictors. According to findings from our previous study (Zhang et al., 2020), the main predictors for virus discovery at the global scale were GDP-related. This suggests that the patterns of virus discovery we have identified may have been largely driven by research effort rather than the underlying biology. In this study, by focusing on more restricted and homogenous regions where the research effort is less variable, we expected to identify predictors more associated with virus biology.

## Materials and Methods

### Data sets of human-infective RNA viruses in three regions

We performed an ecological study, and the subject of interest is each human-infective RNA virus species. With reference to a full list of human-infective RNA virus species (Zhang et al., 2020), we geocoded the first report of each in humans in the United States, China, and Africa separately. The latest version as of 31 December 2019 included 223 species (**Table 1S**), with *Human torovirus* abolished and a new species—*Heartland banyangvirus*—added by ICTV in 2018 (International Committee on Taxonomy of Viruses, 2018). We defined discovery location as where the initial human was exposed to/infected with the virus, as suggested in the first report of human infections from peer-reviewed literature.

We followed the same search terms, databases searched, and inclusion or exclusion criteria as our global data set for data collection (Woolhouse et al., 2018). Reference databases included PubMed, Web of Science, Google Scholar, and Scopus. Two Chinese database [i.e. China National Knowledge Infrastructure (CNKI) and Wanfang Data] were also searched when collecting data for China. Reference lists of relevant studies and reviews were also checked manually to find potential earlier discovery papers. The following key words were used for the retrieval: virus full name or abbreviations or virus synonyms; and human* or person* or case* or patient* or worker* or infection* or disease* or outbreak* or epidemic*; and region name (Chin* or Taiwan or Hong Kong or Macau; United States or US or USA or America*; Africa* or all African country names). Virus synonyms and abbreviations include early names used in the discovery paper and all subtypes provided by the ICTV 10th report (*Full ICTV Online (10th) Report:* https://talk.ictvonline.org/ictv-reports/ictv_online_report/). Evidence which met the following criteria from peer-reviewed literatures were included: (a) Diagnostic methods for RNA virus infection in humans were clearly described, through either viral isolation or serological methods; (b) Specific virus species name or subtypes falling under that species were clearly provided; (c) Both natural infection and iatrogenic or occupational infections were accepted. Evidence which met the following criteria were excluded: (a) Uncertain species due to cross-reactivity with related viruses; (b) Diagnostic methods for virus infection were not specified; (c) Description of clinical symptoms or pathogenicity were not considered as human infection of one certain virus species; (d) Report of ‘[virus name]-like’ or ‘potential [virus name] infections’; (d) Intentional infections including experimental inoculation or vitro infections; (e) Non-peer-reviewed literature, including media reports, thesis, or unpublished data. Literature selection was performed by two individuals independently and discrepancies were resolved by discussion with a third individual.

All locations were geolocated as precisely as possible using methods from our previous paper (Zhang et al., 2020). For each region, a polygon was created for those locations at administrative level 3 (county for the United States; city for China; for Africa, it varies between different countries) and above. Details of data types for virus discovery database in three regions was summarised in **Table 2S**. The majority of discovery locations in the United States and Africa involved point data, while in China the majority involved polygon data at province level. A bootstrap resampling procedure was developed for polygon data covering more than one grid cell (details below). Discovery date of human infection was defined as the publication year in the scientific literature.

### Spatial covariates

As for our global analysis (Zhang et al., 2020), a suite of global gridded climatic, socio- economic, land use, and biodiversity variables (n=33) postulated to affect the spatial distribution of RNA virus discovery were compiled, each at a resolution of 0.5°/30” (except university count having a resolution at country level for Africa and state/province for the United States and China). Data for the United States, China and Africa were extracted by restricting the coordinates within each region. The definition, original resolution, and source of each variable were the same as our previous paper (Zhang et al., 2020). All predictors were aggregated from their original spatial resolution to 1°×1° resolution; data for climatic variables, population, GDP, and land use data without full temporal coverage were extrapolated back to 1901; both following methods from our previous paper (Zhang et al., 2020).

### Boosted regression trees modelling

We used Poisson boosted regression trees (BRT) model to examine the relationship between discovery of RNA virus and 33 predictors for each 1° resolution of grid cell across each region separately, following codes from our previous study (Zhang et al., 2020) and one previous paper (Allen et al., 2017). As a tree-based learning method, BRT model can automatically capture complex relationships and interactions between variables, and also can well account for spatial autocorrelation within the data (Crase et al., 2012). We compared Moran’s I values of the raw virus data and the model residuals to estimate the ability of the BRT model to account for spatial autocorrelation (Cliff et al., 1981). In order to minimise the effect of spatial uncertainty of virus discovery data, we performed 1000 times bootstrap resampling for those discovery locations reported as polygons. We assumed each grid cell in the polygon has the equal chance to be selected, and for each virus record we selected one grid cell randomly from the polygon for each subsample. A ratio of 1:2 for presence to absence constituted each subsample, i.e., for each grid cell with virus discovery, two grid cells with no discovery were randomly selected from ‘virus discovery free’ areas at all time points within the region. Take the United States as an example, each subsample included 95 grid cells with virus discovery and 190 with no virus discovery. We then matched the virus data with all predictors (using the nearest decade for time-varying predictors). We assumed that the virus count in any given grid cell in each decade followed a Poisson distribution, and we calculated the virus discovery count in each grid cell by decade as the response variable.

All BRT models were fitted in R v. 3.6.3, using packages dismo and gbm. BRT models require the user to balance three parameters including tree complexity, learning rate, and bag fraction. Tree complexity reflects the order of interaction in a tree; learning rate shrinks the contribution of each tree to the growing model; bag fraction specifies the proportion of data drawn from the full training data at each step. We set these parameters as recommended from Elith et al (Elith et al., 2008), and make sure each resampling model contained at least 1000 trees. BRT models identified the final optimal number of trees in each model using a 10-fold cross validation stagewise function (Elith et al., 2008). The three parameter values of the optimal model as well as the mean optimal number of trees across 1000 replicate models for all three regions were summarised in **Table 3S**.

By fitting 1000 replicate BRT models, the relative contribution plots and partial dependence plots with 95% quantiles were plotted. We defined variables with a relative contribution greater than the mean (3.03%) as influential predictors in all three regions (Shearer et al., 2018). The partial dependence plots depict the influence of each variable on the response while controlling for the average effects of all the other variables in the model. The map of virus discovery probability across each region in 2010–2019 was derived from the means of the predictions of 1000 replicate models, using values of the 33 predictors in 2015. In order to show discovery hotspots, we converted the prediction map of virus count to a map of probability.

Two statistics were calculated to evaluate the model’s predictive performance: a) the deviance of the bootstrap model (Elith et al., 2008), b) infraclass correlation coefficient (ICC) calculated from 50 rounds of ten-fold cross-validation, by following methods from our previous paper (Zhang et al., 2020). For the ten-fold cross-validation, we selected 50 data sets randomly from the 1000 bootstrapped subsamples. We took the first data set and divided into ten subsets. For each round of ten-fold cross-validation, the unique combinations of 9 subsets constituted the training sets and were used to fit models, and the remaining one was used as a test set to evaluate the predictive performance of the model. We repeat the same process as above for the remaining 49 data sets. One intraclass correlation coefficient (ICC) was calculated from each round of validation and the median with 95% quantiles across all 50 rounds was calculated. The ICC varies between 0 and 1, with an ICC of less than 0.40 representing a poor model, 0.40–0.59 representing a fair model, 0.60–0.74 representing a good model, and 0.75–1 representing an excellent model (Cicchetti, 1994).

Exploratory subgroup analyses (distinguishing viruses firstly discovered in regions and those that had been discovered elsewhere in the world) were performed. We used the same BRT modelling approach as we described above, and relative contribution of each predictor was calculated for each subgroup. We were unable to perform subgroup analysis for China because only 9 human-infective RNA viruses have been firstly discovered in it, and BRT model cannot be fitted to a sample as small as 9.

The R software, version 3.6.3 (R Foundation for Statistical Computing, Vienna, Austria) was used for all statistical analyses. All maps were visualised by using ArcGIS Desktop 10.5.1 (Environmental Systems Research Institute).

## Results

The numbers of human-infective virus species discovered in the United States, China and Africa up to October 2019 were 95, 80 and 107 respectively (**Table 1S**). Most first discoveries have been in eastern United States (especially in areas around Maryland, Washington, D.C., and New York), eastern China (developed cities including Beijing, Hong Kong, Shanghai, and Guangzhou), and southern and central Africa (Pretoria and Johannesburg, South Africa; Borno State and Ibadan, Nigeria) (**Figure 1**). A total of 60 virus species were previously reported in all three regions, and 27, 12, 37 species were only found in the United States, China, and Africa respectively (**Figure 2**). In all three regions, smaller proportions of viruses were vector-borne [United States: 23.2% (22/95); China: 21.3% (17/80); Africa: 27.1% (29/107)] and strictly zoonotic [United States: 30.5% (29/95); China: 16.3% (13/80); Africa: 33.6% (36/107)], compared to large proportions for both virus types at the global scale [vector-borne: 41.7% (93/223) and strictly zoonotic: 58.7% (131/223)] (**Figure 2**). The 60 shared species were also disproportionally vector-borne [11.7% (7/60)] and strictly zoonotic [7% (4/60), **Figure 2**].

**Figure 1.**
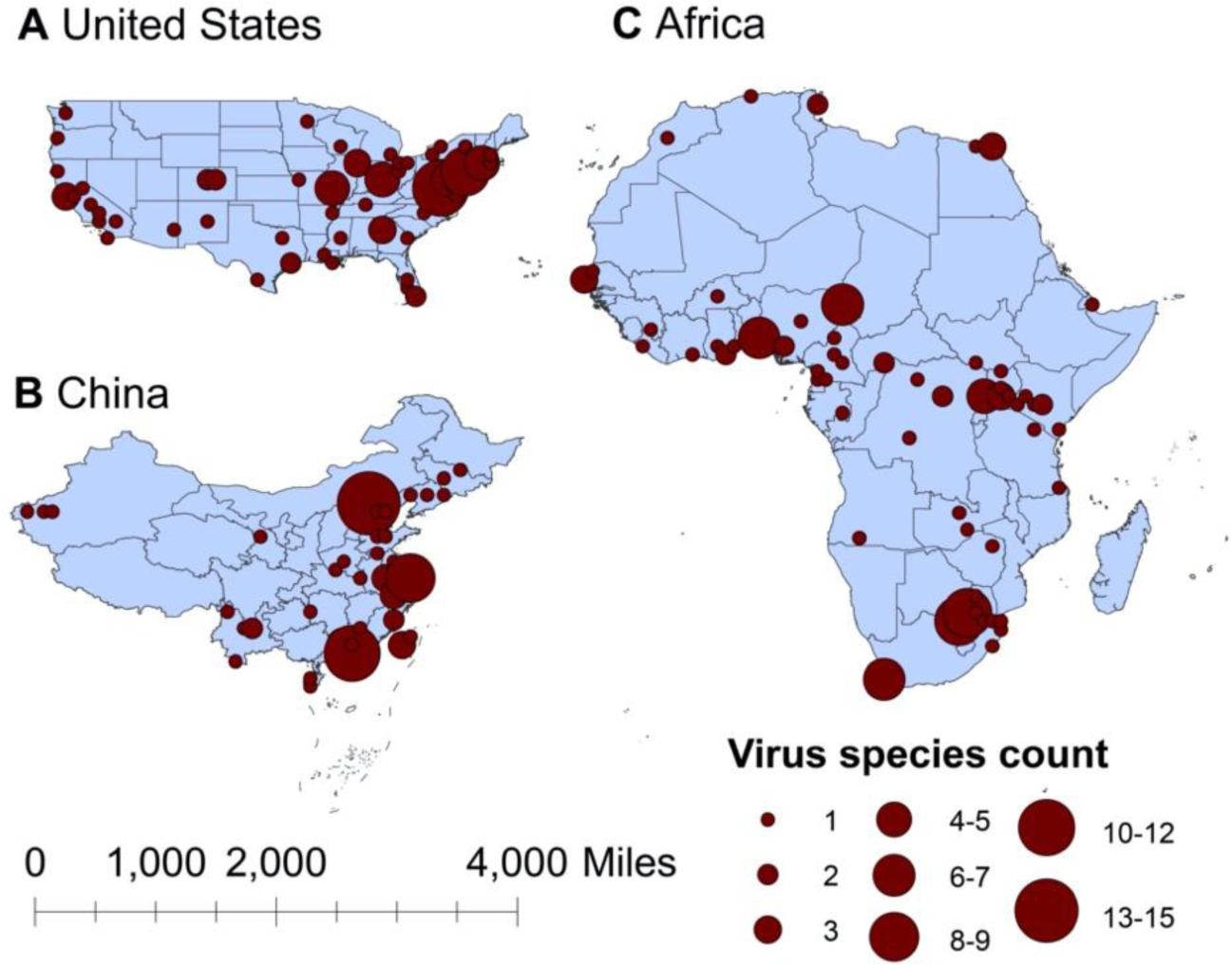
Spatial distribution of human-infective RNA virus discovery in three regions, 1901–2019. A, United States. B, China. C, Africa. Red dots represent discovery points or centroids of polygons, with the size representing the cumulative virus species count.

**Figure 2.**
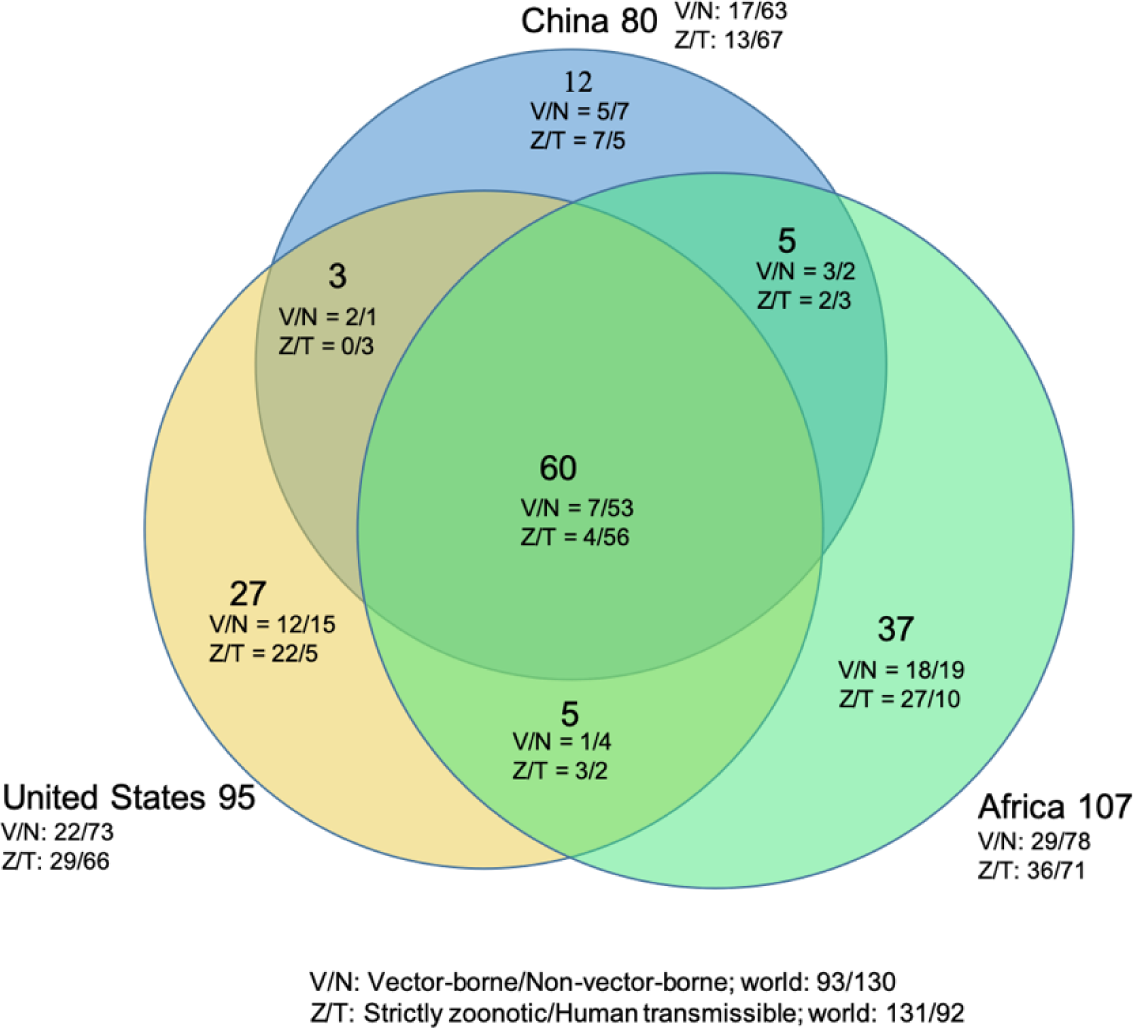
Shared human-infective RNA virus species count in three regions. Under/By the species count the ratios of vector-borne (V) to non-vector-borne (N) viruses and strictly zoonotic (Z) to human transmissible (T) viruses were shown.

The discovery curves for the United States and Africa have seen a broadly similar pattern, with China lagging behind these two regions (**Figure 3**). In comparison with the world, the median time lag of the virus discovery was 0 [interquartile range (IQR): 2.5], 12 (IQR: 29.5) and 2 (IQR: 10.5) years in the United States, China and Africa, respectively (**Figure 1S**). In China, the time lag was noticeably shorter for viruses discovered after 1975 [before 1975: a median lag of 30.5 (IQR: 30.5) years; after 1975: 2.5 (IQR: 7) years, p value of Wilcoxon rank sum test < 0.001].

**Figure 3.**
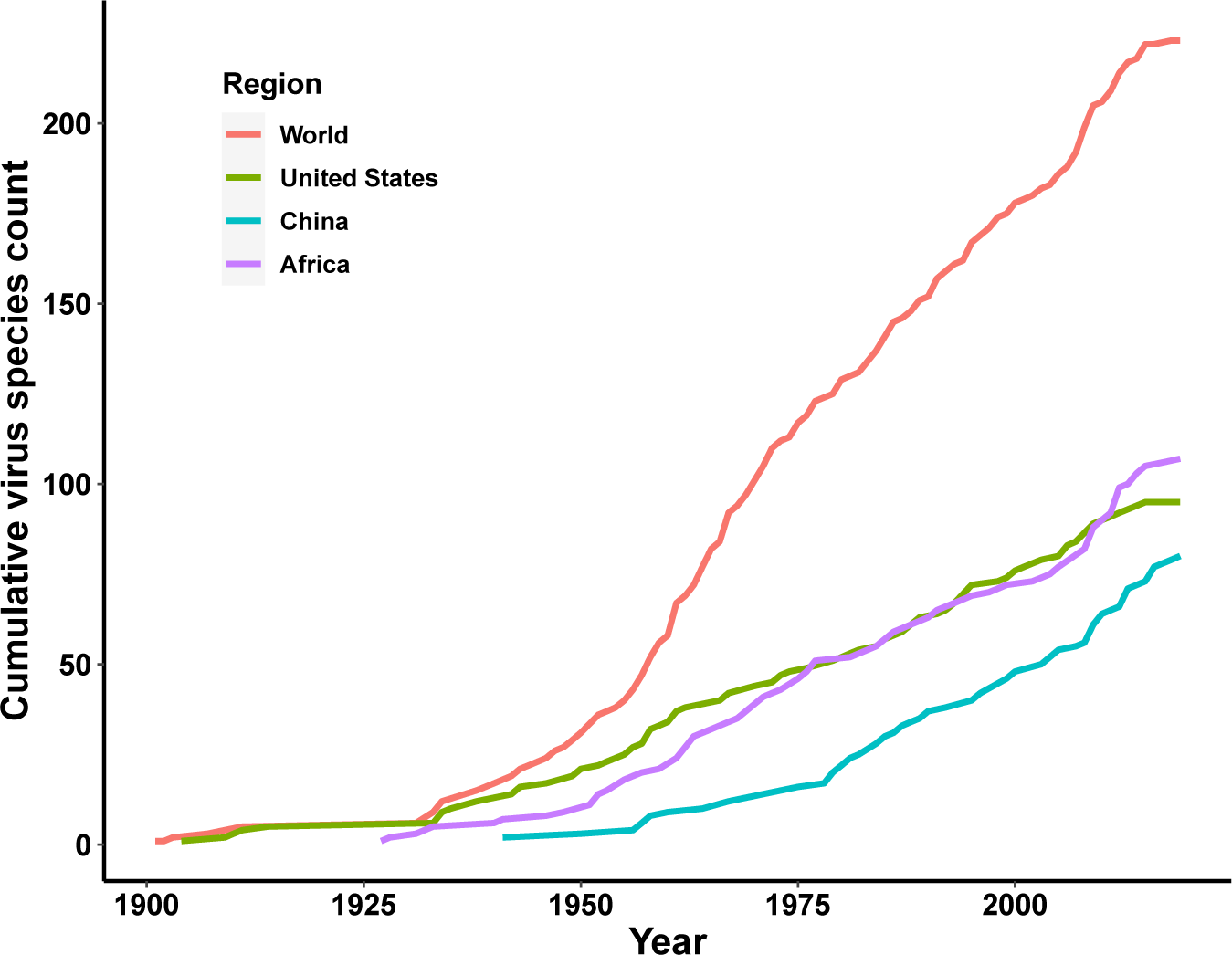
Discovery curve of human-infective RNA virus species in three regions and the world

In the United States, six variables including three predictors related to land use [urbanized land: relative contribution of 35.8%, urbanization of cropland (i.e. the percentage of land area change from cropland to urban land): 8.0%, growth of urbanized land: 4.1%], two socio- economic variables (GDP growth: 10.0%; GDP: 5.7%), and one climatic variable (diurnal temperature change: 4.9%) were identified as important predictors for discriminating between locations with and without virus discovery (**Figure 4A**). The partial dependence plots shown in **Figure 2S** suggested non-linear relationships between the probability of virus discovery and most predictors. All important predictors presented a positive trend over narrow ranges at lower values.

**Figure 4.**
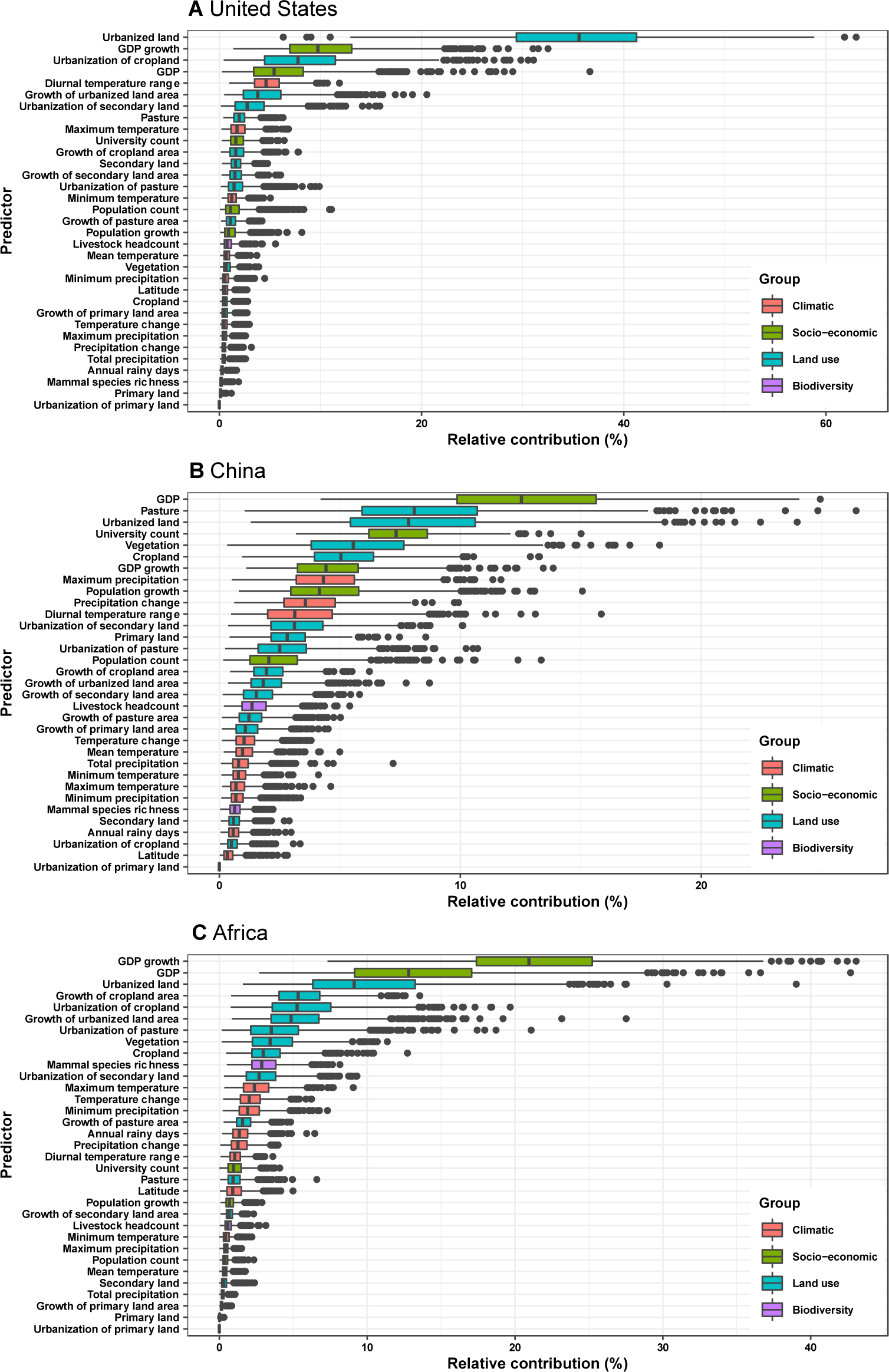
Relative contribution of predictors to human-infective RNA virus discovery in three regions. A, United States. B, China. C, Africa. The boxplots show the median (black bar) and interquartile range (box) of the relative contribution across 1000 replicate models, with whiskers indicating minimum and maximum and black dots indicating outliers.

In China, twelve variables including four socio-economic variables (GDP: 12.7%, university count: 7.5%, GDP growth: 4.6%, population growth: 4.4%), five predictors involving land use [pasture: 8.3%, urbanized land: 8.1%, vegetation: 5.8%, cropland: 5.3%, urbanization of secondary land (the percentage of land area change from secondary land to urban land; secondary land is natural vegetation that is recovering from previous human disturbance): 3.3%], and three climatic variables (maximum precipitation: 4.5%, precipitation change: 3.8%, diurnal temperature range: 3.3%) were identified as important predictors for discriminating between locations with and without virus discovery (**Figure 4B**). GDP, urbanized land, university count, vegetation, GDP growth, maximum precipitation, population growth, and urbanization of secondary land presented a positive trend over narrow ranges at lower levels; pasture, cropland, precipitation change, and diurnal temperature range had non-monotonic/ negative impacts, with highest risks at lower values (**Figure 3S**).

In Africa, ten variables including two socio-economic variables (GDP growth: 21.2%, GDP: 13.0%), seven predictors related to land use (urbanized land: 9.4%, growth of cropland area: 5.6%, urbanization of cropland: 5.5%, growth of urbanized land: 5.1%, urbanization of pasture: 3.8%, vegetation, 3.7%, cropland: 3.2%), and one biodiversity variable (mammal species richness: 3.1%) were identified as important predictors for discriminating between locations with and without virus discovery (**Figure 4C**). All important predictors presented a positive trend over narrow ranges at lower positive values, except mammal species over a large range (**Figure 4S**).

Our BRT models reduced Moran’s I value below 0.15 in all three regions (**Figure 5S**), suggesting that BRT models with 33 predictors have adequately accounted for spatial autocorrelations in the raw virus data in all three regions. The model validation statistics for each region are shown in **Table 4S**. Combining these measures, our BRT model predictions range from fair to good (Cicchetti, 1994).

**Figure 5.**
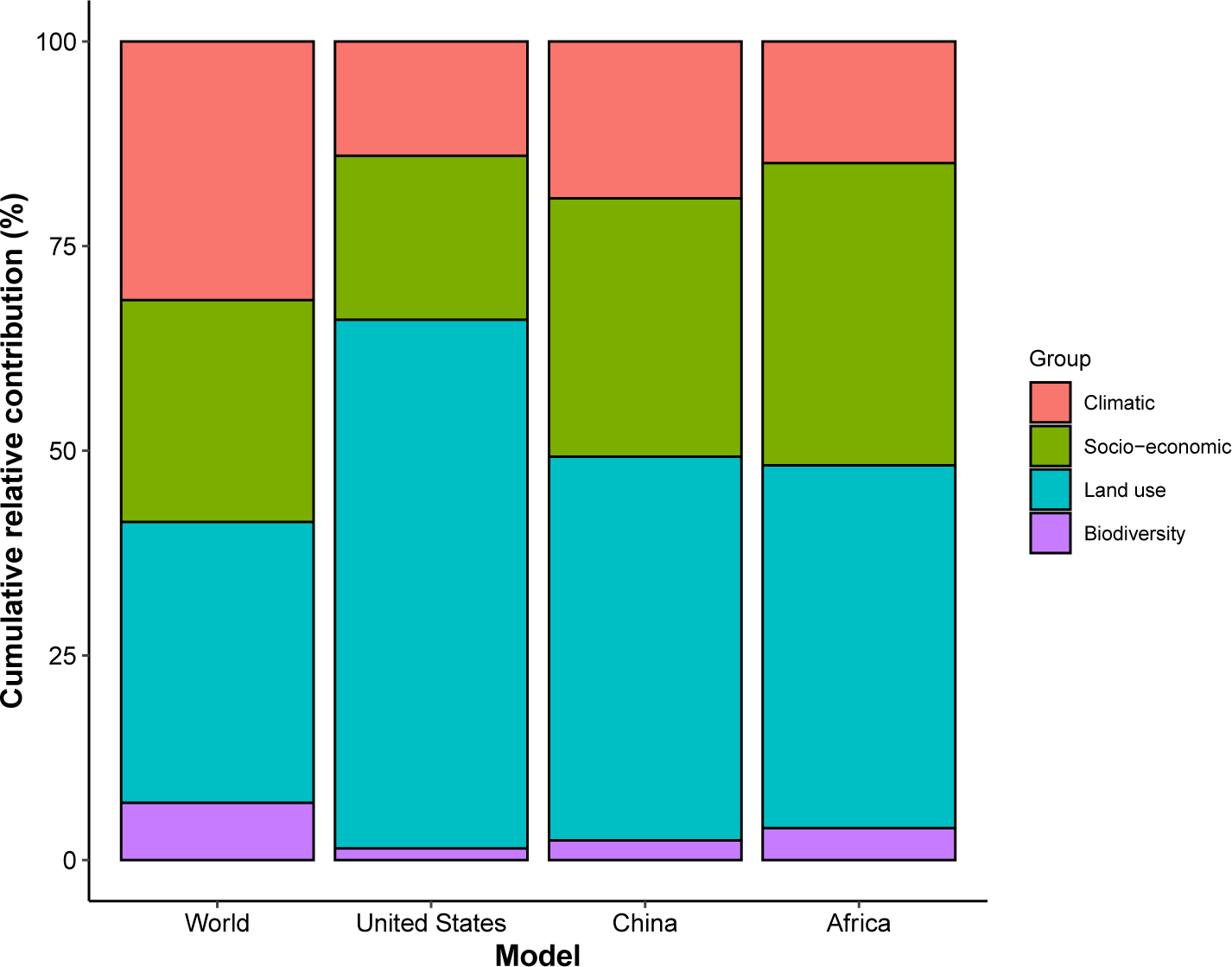
Cumulative relative contribution of predictors to human-infective RNA virus discovery by group in each model of different regions. The relative contributions of all explanatory factors sum to 100% in each model, and each colour represents the cumulative relative contribution of all explanatory factors within each group.

In all three regions, human-infective RNA virus discovery was best predicted by land use and socio-economic variables, followed by climatic variables and biodiversity (**Figure 5**), whereas virus discovery was more associated with climatic variables and biodiversity at the global level. The comparison between three regions showed that: climatic variables contributed most to the discovery of human-infective RNA viruses in China; land use contributed most to the discovery in the United States; socio-economic variables and biodiversity contributed most to the discovery in Africa and least to discovery in the United States.

We mapped human-infective RNA virus discovery probability in 2010–2019 for the three regions, based on the fitted BRT models and values of all 33 predictors in 2015 (**Figure 6S**– **Figure 8S**). Outside contemporary risk areas where human-infective RNA viruses were previously discovered in the United States (**Figure 1A**), we predicted high probabilities of virus discovery across southern Michigan, central-Northern Carolina, central Oklahoma, southern Nevada, and north-eastern Utah (**Figure 6A**). Outside contemporary risk areas where human-infective RNA viruses were previously discovered in China (**Figure 1B**), we predicted high probabilities of virus discovery across other eastern China area as well as two western areas including south-central Shaanxi and north-eastern Sichuan (**Figure 6B**). Outside contemporary risk areas where human-infective RNA viruses were previously discovered in Africa (**Figure 1C**), we predicted high probabilities of virus discovery across northern Morocco, northern Algeria, northern Libya, south-eastern Sudan, central Ethiopia and western Democratic Republic of the Congo (**Figure 6C**). Most new virus species since 2010 in each region (6/6 in the United States, 19/19 in China, 12/19 in Africa) were discovered in high-risk areas (85% percentiles of predicted probability across each region) as predicted by our model. Of all the 37 (United States: 6; China: 19; Africa: 12) viruses discovered in high-risk areas in 2010–2019, 13 (United States: 2; China: 7; Africa: 4) viruses were discovered at the potential new hotspots where there have not been any virus discoveries before 2010.

**Figure 6.**
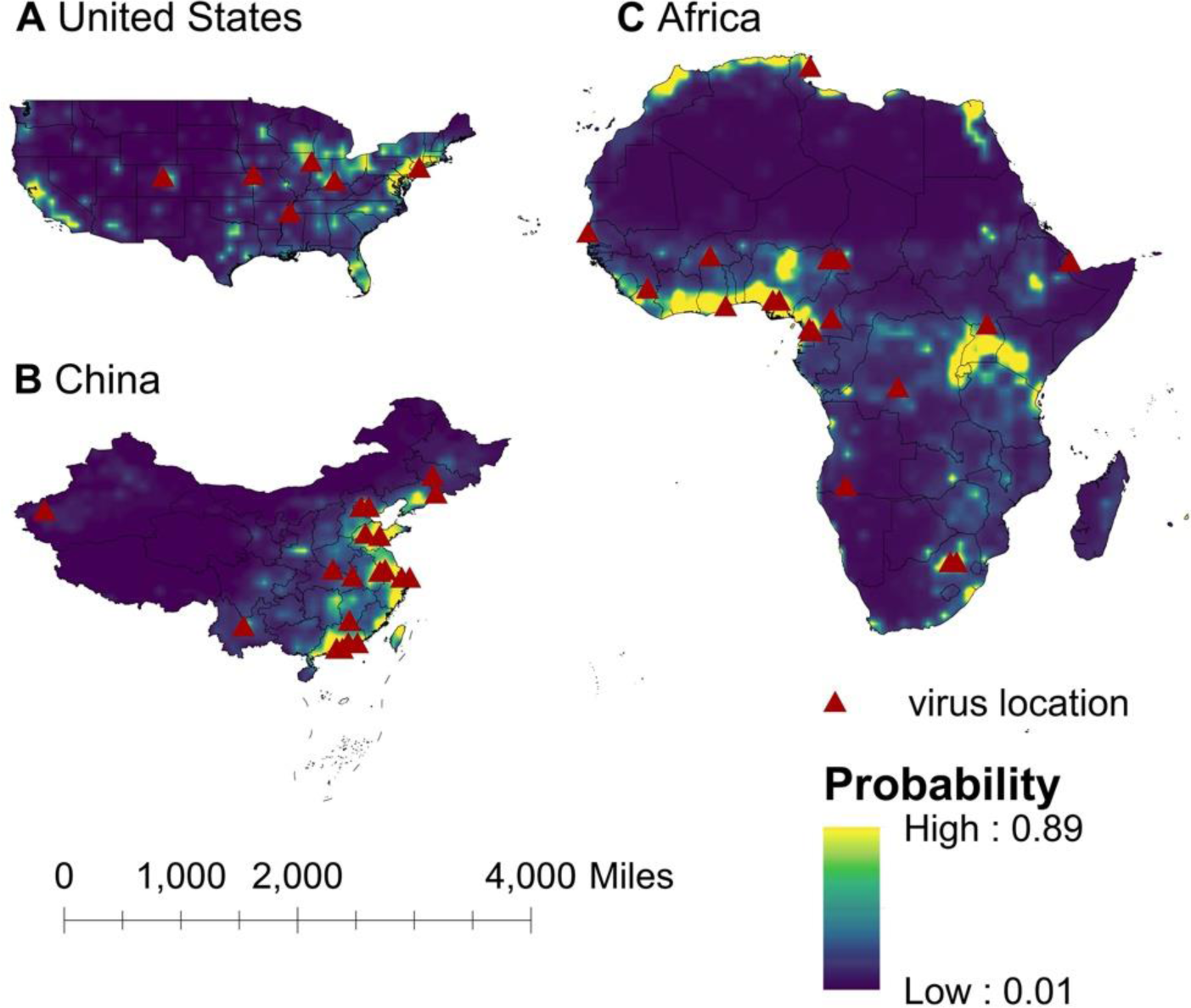
Predicted probability of human-infective RNA virus discovery in three regions in 2010–2019. A, United States. B, China. C, Africa. The triangles represented the actual discovery sites from 2010 to 2019, and the background colour represented the predicted discovery probability.

Based on our subgroup analysis (distinguishing viruses firstly discovered in regions and those that had been discovered elsewhere in the world), discoveries of human-infective RNA viruses firstly discovered from either United States or Africa were better predicted by climatic and biodiversity variables, while discoveries of viruses that had been discovered from elsewhere in the world were better predicted by socio-economic variables (**Figure 9S**).

## Discussion

To our knowledge, this analysis represents the first investigation of human-infective RNA virus discovery in three large regions of the world which have experienced distinct socio- economic, ecological and environmental changes over the last 100 years. In total, 95 human-infective RNA virus species had been found in the United States; 80 in China; 107 in Africa.

The discovery maps of human-infective RNA virus in the three regions indicated areas with historically high discovery counts: eastern and western United States, eastern China, and central and southern Africa. BRT modelling suggested that the relative contribution of 33 predictors to human-infective RNA virus discovery varied across three regions, though climatic and biodiversity variables were consistently less important in all three regions than at a global scale. We mapped the probability of human-infective RNA virus discovery in 2010– 2019 which would continue to be high in historical hotspots but, in addition, we identified several new hotspots in central-eastern and southwestern United States, eastern and western China, and northern Africa. These results offer a tool for public health practitioners and policy-makers to better understand local patterns of virus discovery and to invest efficiently in surveillance systems at the local level.

In all three regions, GDP and/or GDP growth were identified as important predictors for virus discovery, especially in Africa where GDP and GDP growth were identified as the leading predictors. This is consistent with our previous analysis that GDP and GDP growth play a major role in discovering viruses (Zhang et al., 2020). In general, sufficient economic, human and material resources, the availability of advanced infrastructure and technology, and greater research capabilities in the relative higher-income areas enable the virus discovery (Rosenberg et al., 2013). That this effect applied both within one continent and within single countries such as the United States and China suggested that most virus discoveries were likely passive, i.e. the viruses were detected when they arrived in a location with the resources to detect them. This is plausible because in all regions in our study, human-transmissible viruses accounted for the larger proportion, and our previous analysis suggested richer areas were more likely to first capture transmissible viruses (e.g. Influenza virus, Rhinovirus, Rabies lyssavirus, Measles morbillivirus, Mumps orthorubulavirus, Rubella virus, and Norwalk virus) capable of spreading to multiple areas (Zhang et al., 2020). Temporally, in China the rate of discovery increased after economic growth accelerated in the 1980s (**Figure 3**). We note in publications describing first virus discoveries that most historical virus discoveries in Africa received support from the United States and Europe, and this may explain why Africa saw an increased number of virus discoveries after 1950—30 years earlier than China (**Figure 3**). Notably, in China the relative contribution of GDP growth to virus discovery was not as substantial as that in Africa. In contrast, university count was found to be associated with virus discovery, suggesting virus discovery likely being a significant area of research in Chinese universities. Our model also suggested the overall socio-economic factors contributed less in the United States than other two regions. The possible explanation is that the socio-economic level across the whole United States is relatively high and homogenous.

Predictors other than GDP and university count are likely to be linked to virus natural history. In all three regions, the area of urban land and further urbanization made great contribution to virus discovery. This reinforced previous studies that urbanization was linked to the detection of new human pathogens through the denser urban population, increased human-wildlife contact rate, spill-over of human infection from enzootic cycle, and the contamination of the urban environment with microbial agents (Hassell et al., 2017; Olival et al., 2017; Weaver, 2013). In the United States, land use contributed most to virus discovery in comparison to other regions—urbanized land, urbanization of cropland, and growth of urbanized land alone had a relative contribution of 47.9%. However, the leading role of urbanization to virus discovery in the United States remains unclear. There might be many factors contributing to this, e.g. the underlying relationships between urbanization and economic growth as well as population growth, but we can’t untangle these from this study.

Consistent with our previous work, population growth was identified as a less prominent predictors (Zhang et al., 2020). In China, population growth—though with greater influence than other regions—contributed less than urbanized land and three other land types on virus discoveries. This reinforces our previous interpretation that urbanization brings larger changes on human living environment than human population size/growth, and therefore may have influenced virus discovery more greatly.

Climate had less influence on human-infective RNA virus discovery in all three regions in comparison to other predictors, in contrast to virus discovery at a global scale (Zhang et al., 2020). The underlying reason may be that the proportion of vector-borne viruses—whose distribution and abundance is strongly associated with the impact of climate on vector populations (Li et al., 2014)—in all three regions (United States: 23.2%; China: 21.3%; Africa: 27.1%) were less than that in the world (41.7%) (**Figure 3**). Vector-borne viruses tend to have more restricted global ranges, so are less likely to appear in a study of any one region (Zhang et al., 2020).

In addition, a relative smaller proportion of strictly zoonotic viruses in three regions (United States: 30.5%; China: 16.3%; Africa: 33.6%) than that in the world (58.7%) (**Figure 2**) made biodiversity contribute less to virus discovery in the three regions than in the world (Zhang et al., 2020). With exposure to denser mammals played a slightly larger role in virus discovery in Africa than in China and the United States (**Figure 6S–Figure 8S**).

Our discovery probability maps for 2010–2019 in three regions captured most historical hotspots, though several small new areas in central-eastern and southwestern United States, eastern and western China, as well as northern Africa would also make greater contribution to virus discovery (**Figure 6**). Our model has a good predictive ability, given 84% (37/44) new virus species in 2010–2019 were discovered in high-risk areas we have defined—85% percentiles of discovery probability within each region. Further, 35% (13/37) of those viruses discovered in high-risk areas since 2010 were discovered at the potential new hotspots where there had not been any virus discoveries in the past.

Our subgroup analyses suggested in both the United States and Africa, discoveries of viruses firstly discovered in regions were more likely to be associated with climatic and biodiversity variables while discoveries of viruses had been discovered elsewhere in the world were more likely to be associated with socio-economic variables. This is plausible, again because after a novel virus was discovered elsewhere in the world, it is usually areas with a higher socio- economic level firstly capture the virus in the local region.

This study had limitations. First, one common problem for data collected from literature review is the time lag between virus discovery and publication, in which case the virus data are likely to be matched to covariates in later decades. Second, we acknowledge that it is possible we have not identified the earliest report for some well-known viruses such as yellow fever virus, measles virus, especially in the post-vaccination era. Third, we were unable to identify robust and comprehensive data for all three regions on virus discovery effort, although we interpret GDP and university count as being an indirect measure of resources available for this activity.

The study adds to our previous study (Zhang et al., 2020) in several ways. First, we firstly construct data sets of human-infective RNA virus discovery reflecting the viral richness in three broad regions of the world. Second, we reduced the heterogeneity of the predictors by focusing on regions, including those predictors reflecting the research effort. Research effort is less variable within restricted regions and therefore has less effect on virus detection. This implies our predicted hotspots stand closer to the virus geographic distribution in nature. Third, the predicted hotspots derived from regional analysis have a higher precision than at a global scale, e.g. specific areas in the United States and China were identified as hotspots from regional analysis, rather than the whole eastern area from the global analysis. This helps target areas for future surveillance.

In conclusion, a heterogeneous pattern of virus discovery-driver relationships was identified across three regions and the globe. Within regions virus discovery is driven more by land-use and socio-economic variables; climate and biodiversity variables are consistently less important predictors than at a global scale. We mapped with good accuracy that in 2010– 2019 three regions where human-infective RNA viruses had previously been discovered would continue to be the discovery hotspots, but in addition, several new areas in each region would make great contribution to virus discovery. Results from the study could guide active surveillance for new human-infective viruses in high risk areas.

## Author contributions

Feifei Zhang, Conceptualization, Data preparation, Formal analysis, Methodology, Validation, Visualization, Interpretation, Writing - original draft, Writing - review and editing; Margo Chase-Topping, Methodology, Interpretation, Writing - review and editing; Chuan-Guo Guo, Data preparation, Methodology, Validation, Interpretation, Writing - review and editing; Mark E.J. Woolhouse, Conceptualization, Methodology, Interpretation, Writing - original draft, Writing - review and editing, Funding Acquisition.

## Data availability

The authors confirm that all data or the data sources are provided in the paper and its Supplementary Materials. The final datasets and codes used for the analyses are available via figshare at https://doi.org/10.6084/m9.figshare.15101979.

## Supplementary materials

### Supplementary tables

**Table 1S.**
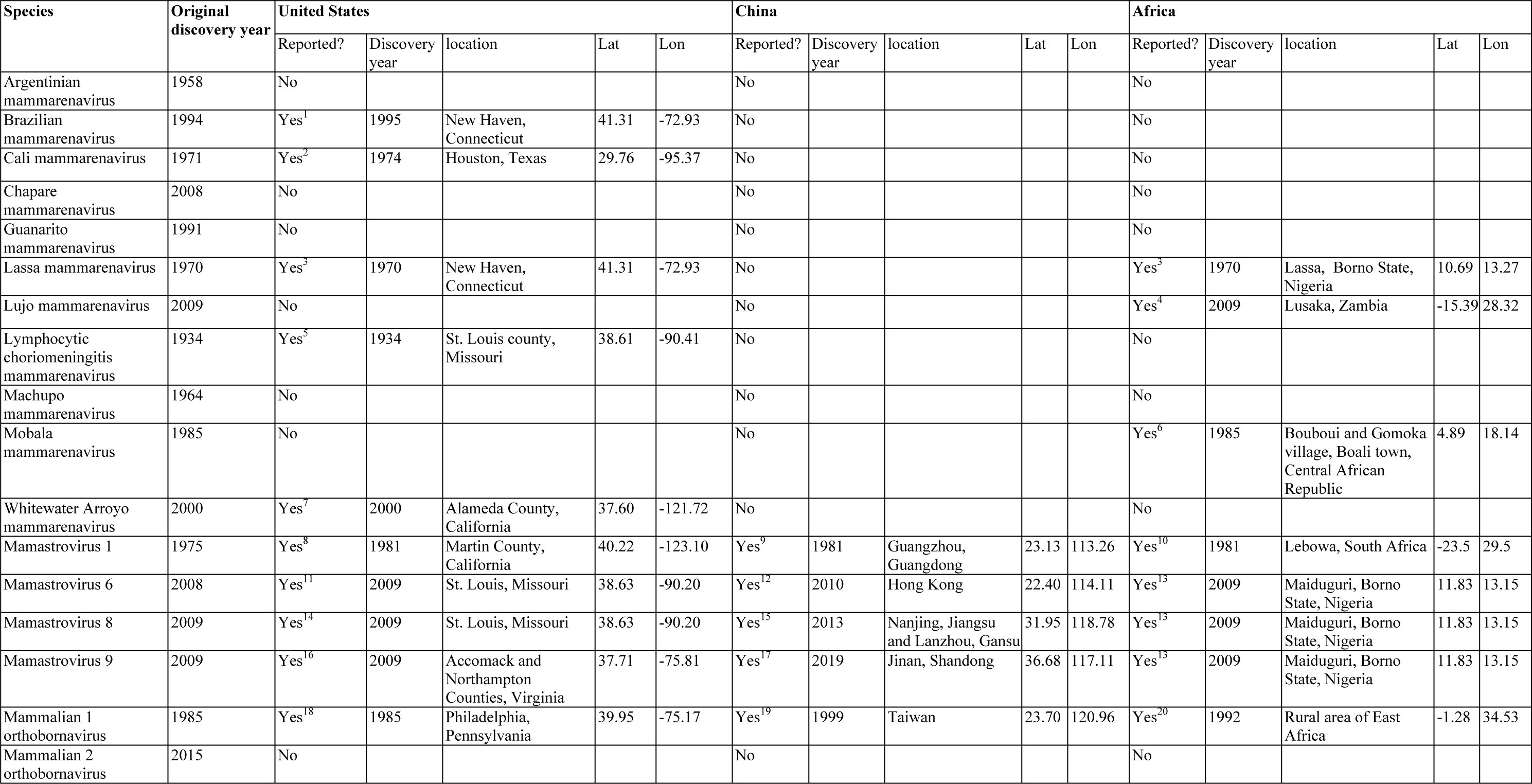
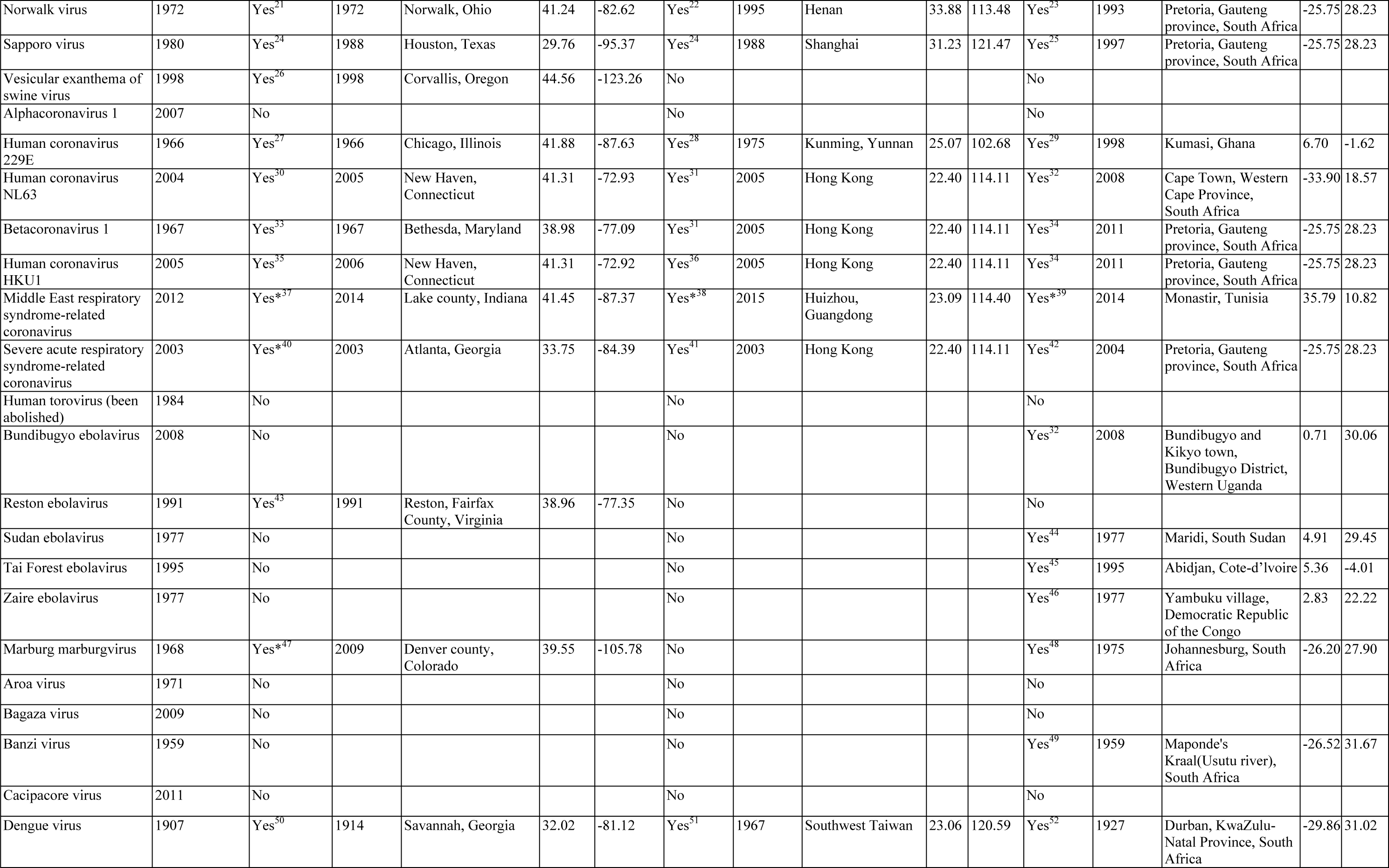
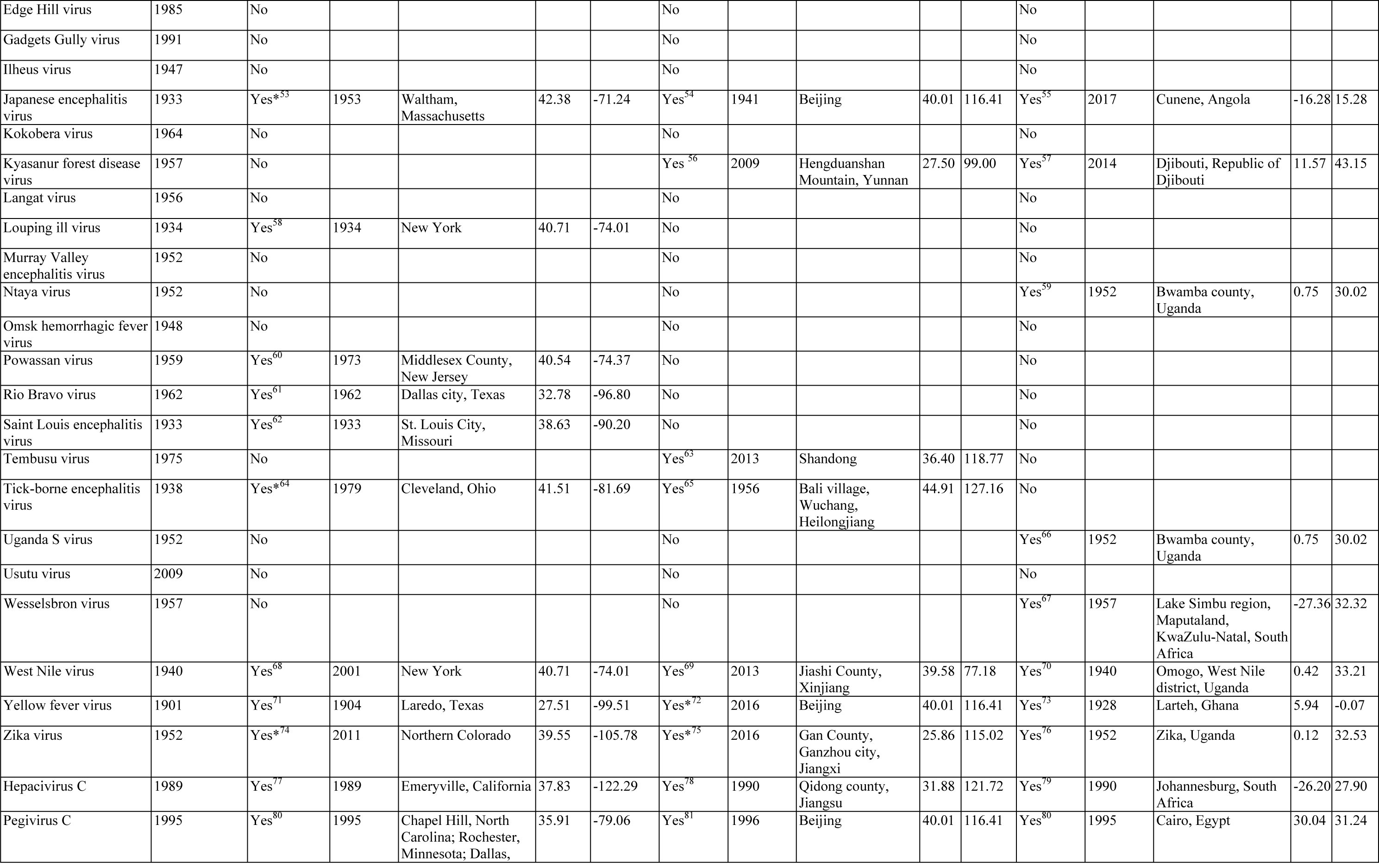
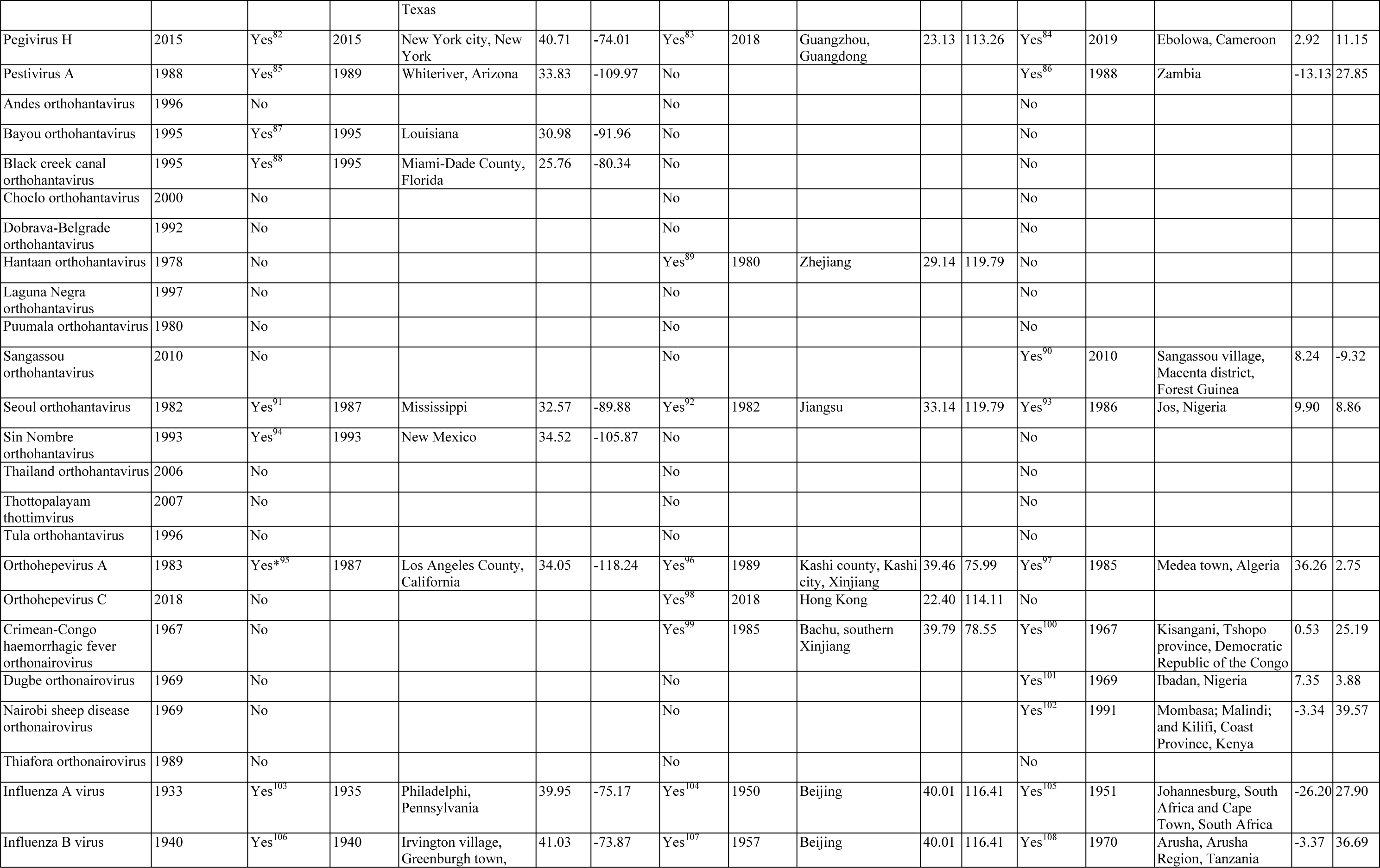
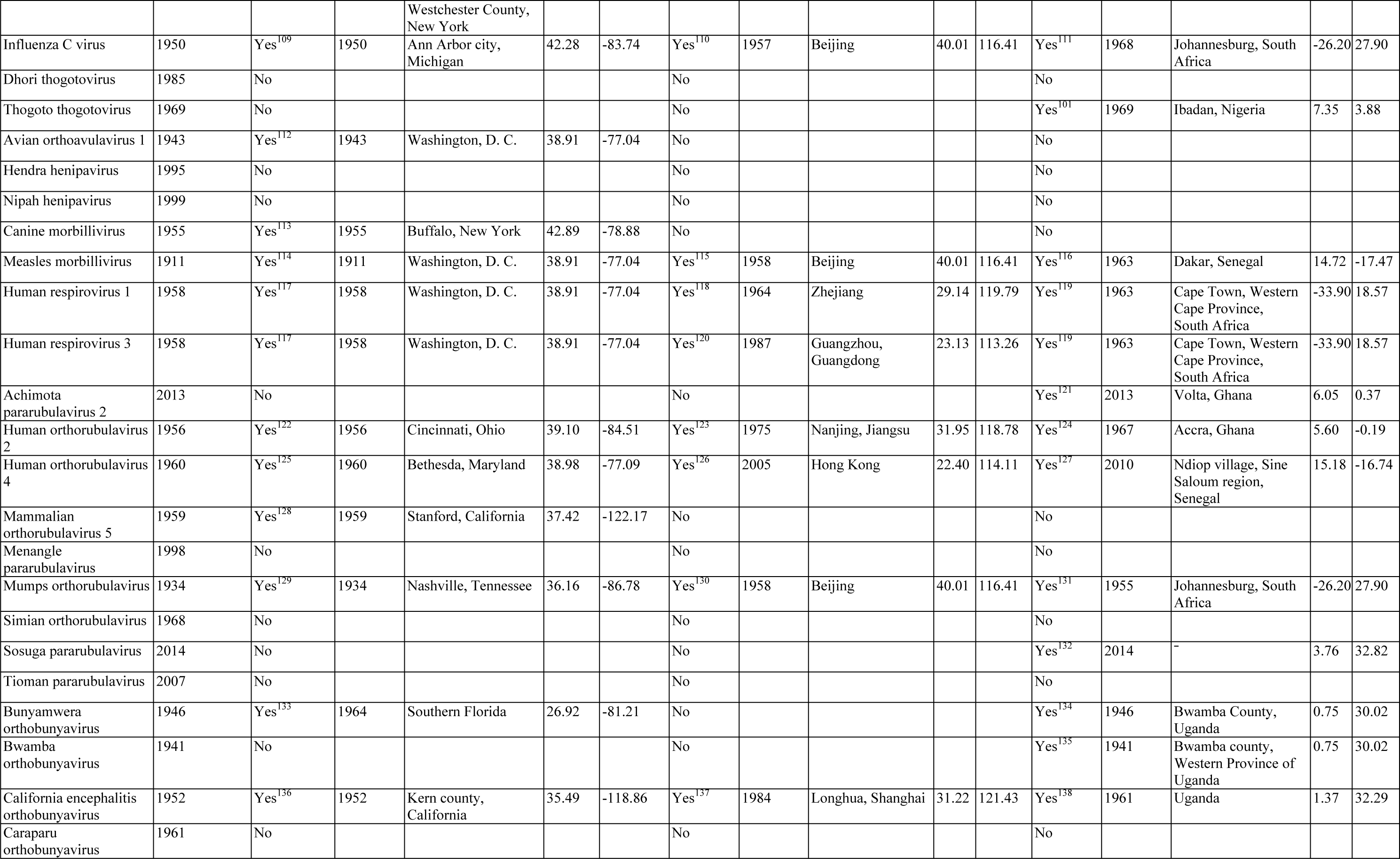
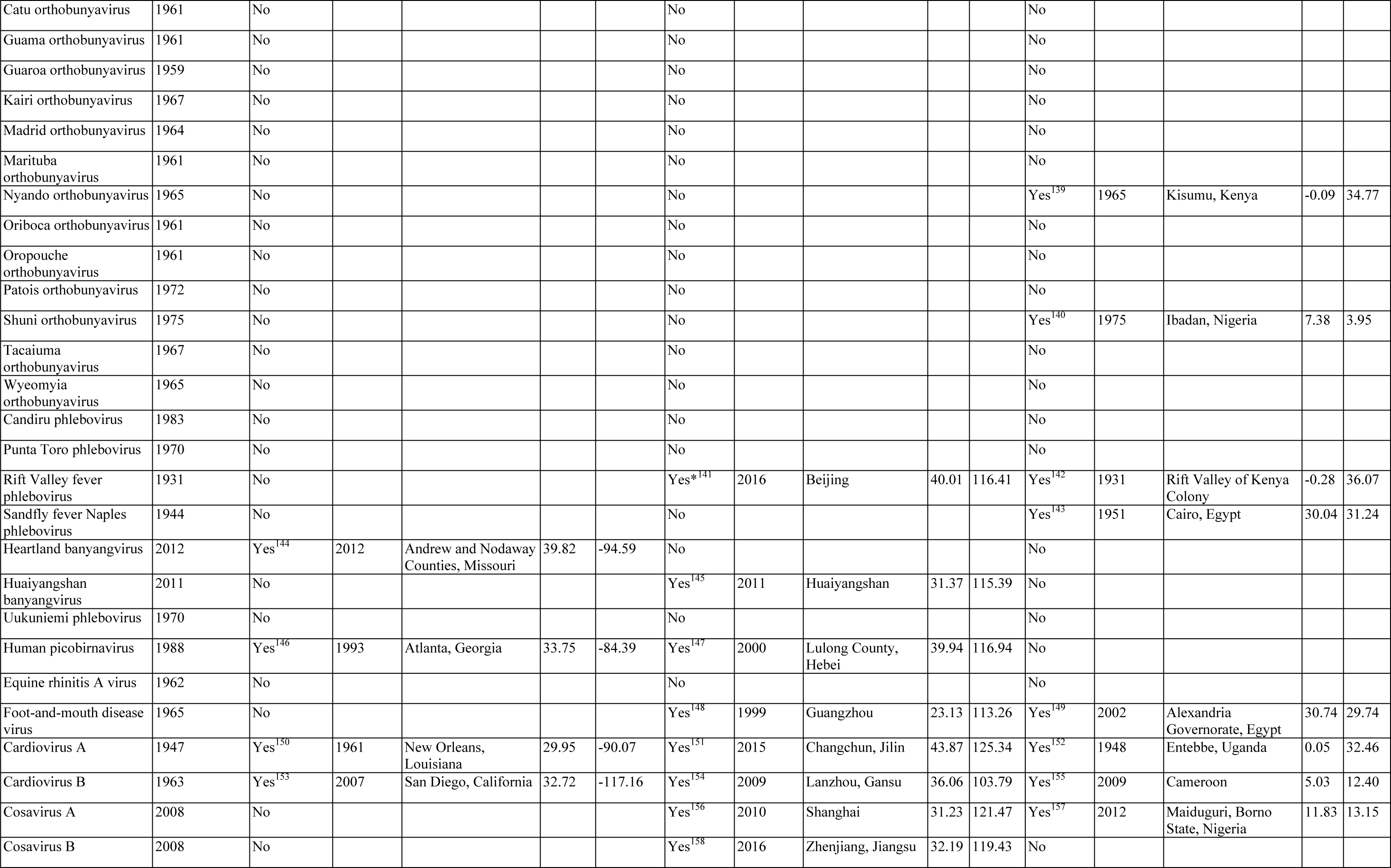
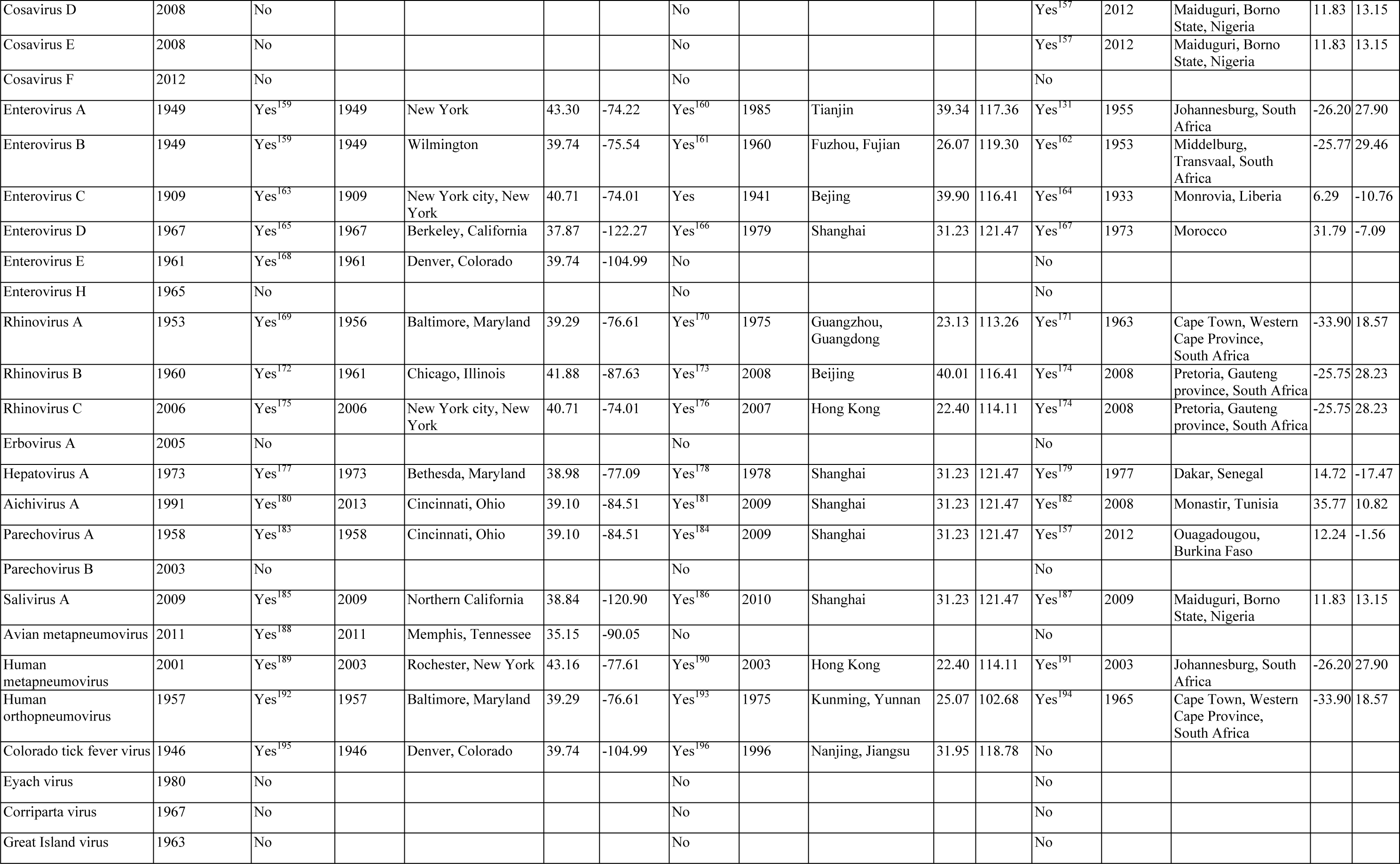
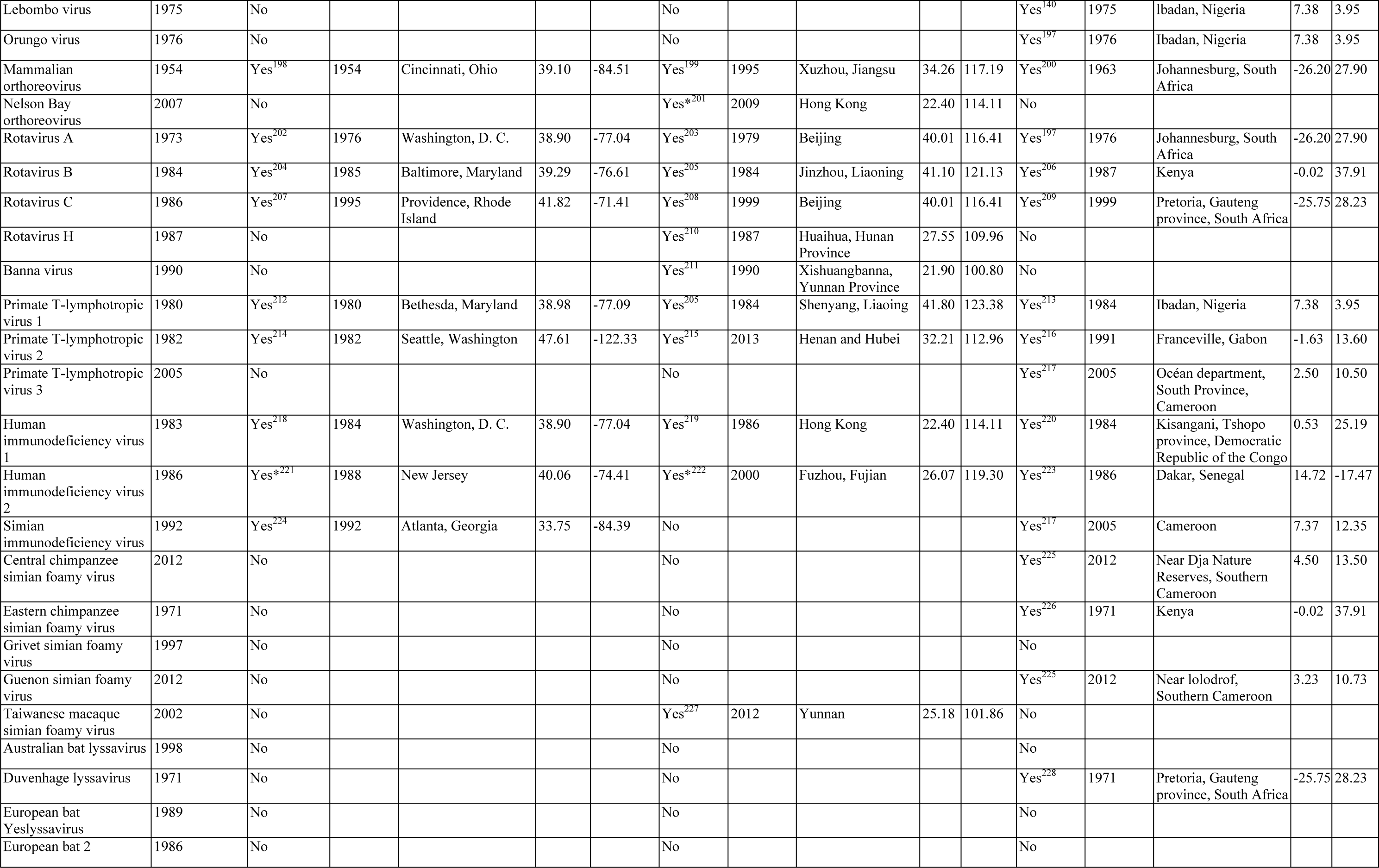
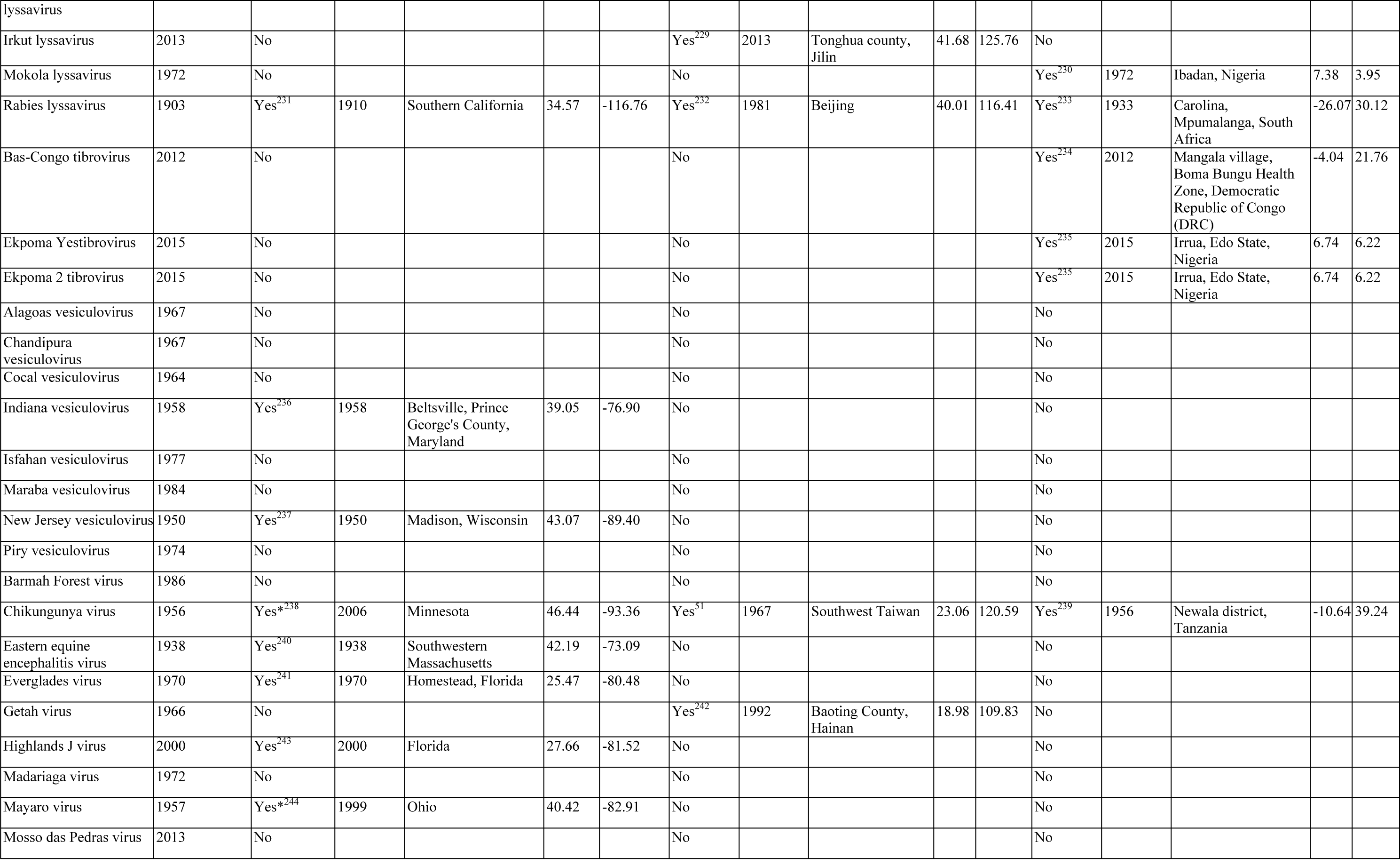
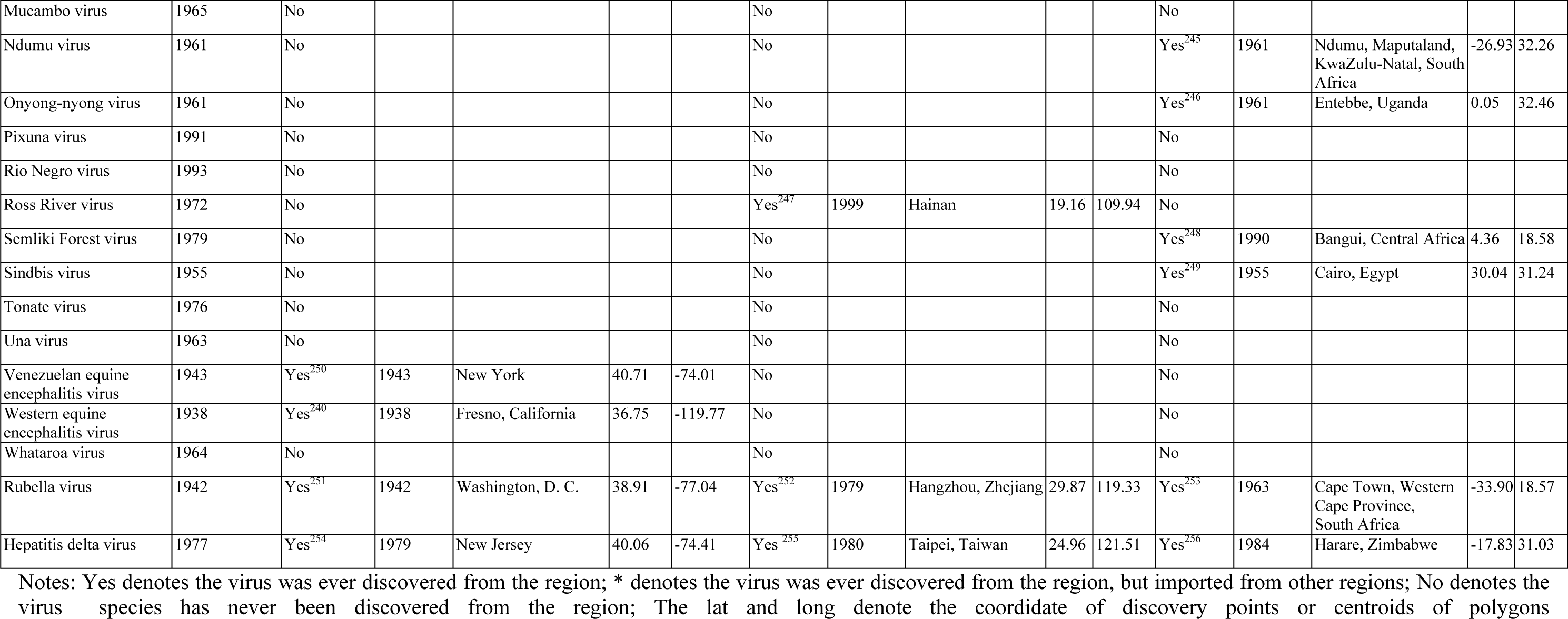
Summary of the human-infective RNA virus data sets in the United States, Africa and China

**Table 2S.** Resolution and covered grid cells for virus discovery data

|  |  | Polygon data |  |  | Point data | Total |
| --- | --- | --- | --- | --- | --- | --- |
|  |  | Country level | State/Province level | City/County level |  |  |
| United States | Virus counts | NA | 14 (14.7%) | 11 (11.6%) | 70 (73.7%) | 95 |
|  | Gridded cell counts | NA | 189 | 12 | 72* | 273 |
| China | Virus counts | NA | 22 (27.5%) | 47 (58.7%) | 11 (13.8%) | 80 |
|  | Gridded cell counts | NA | 161 | 70 | 12* | 243 |
|  | Virus counts | 7 (6.5) | 5 (4.7%) | 15 (14.0%) | 80 (74.8%) | 107 |
| Africa | Gridded cell counts | 307 | 22 | 17 | 80 | 426 |

\*Grid cell counts here include viruses first detected in multiple points from the literature, NA, not applicable

**Table 3S.**
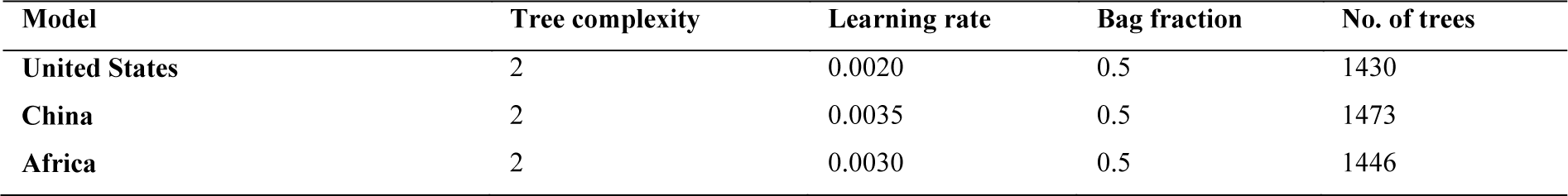
Model parameters

| Model | Tree complexity | Learning rate | Bag fraction | No. of trees |
| --- | --- | --- | --- | --- |
| United States | 2 | 0.0020 | 0.5 | 1430 |
| China | 2 | 0.0035 | 0.5 | 1473 |
| Africa | 2 | 0.0030 | 0.5 | 1446 |

**Table 4S.** **Model validation statistics for analyses in three regions**

| Model | % of deviance explained (95% quantiles) | ICC (95% quantiles) |
| --- | --- | --- |
| United States | 50.5% (44.3%–56.8%) | 0.66 (0.60–0.70) |
| China | 42.0% (32.4%–50.8%) | 0.52 (0.41–0.60) |
| Africa | 42.4% (34.2%–50.0%) | 0.51 (0.44–0.62) |

ICC, intraclass correlation coefficient

### Supplementary Figures

**Figure 1S.**
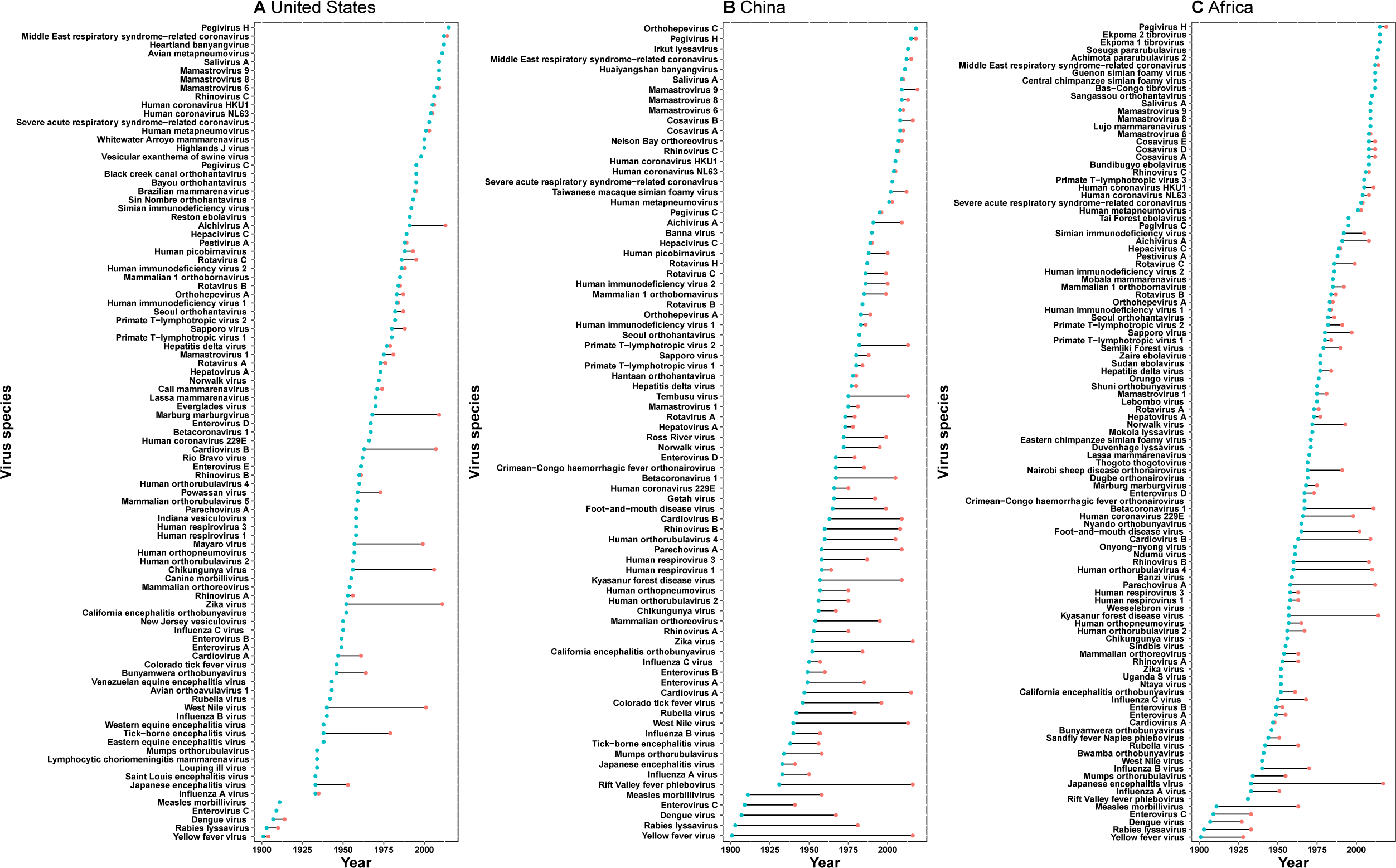
Time lag of human-infective RNA virus discovery between the three regions and the world. A, United States. B, China. C, Africa. The blue dots represent the original discovery year of each virus in the world; the red dots represent the discovery year of each virus in three regions; and the segments between them represent the time lag.

**Figure 2S.**
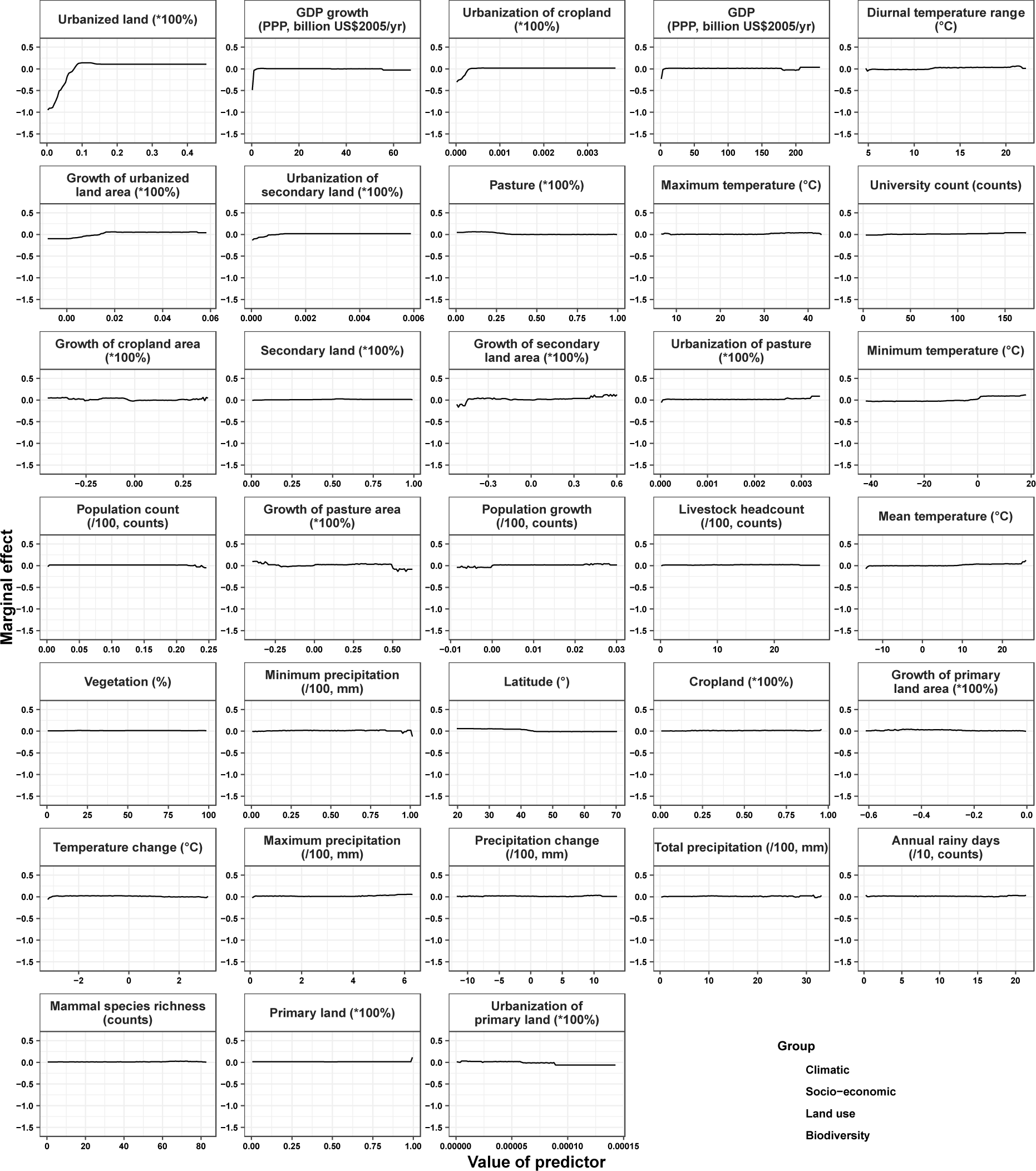
Partial dependence plots showing the influence on human-infective RNA virus discovery for all predictors in the Unites States. Partial dependence plots show the effect of an individual explanatory factor over its range on the response after factoring out other explanatory factors. Fitted lines represent the median (black) and 95% quantiles (coloured) based on 1000 replicated models. Y axes are centred around the mean without scaling. X axes show the range of sampled values of explanatory factors.

**Figure 3S.**
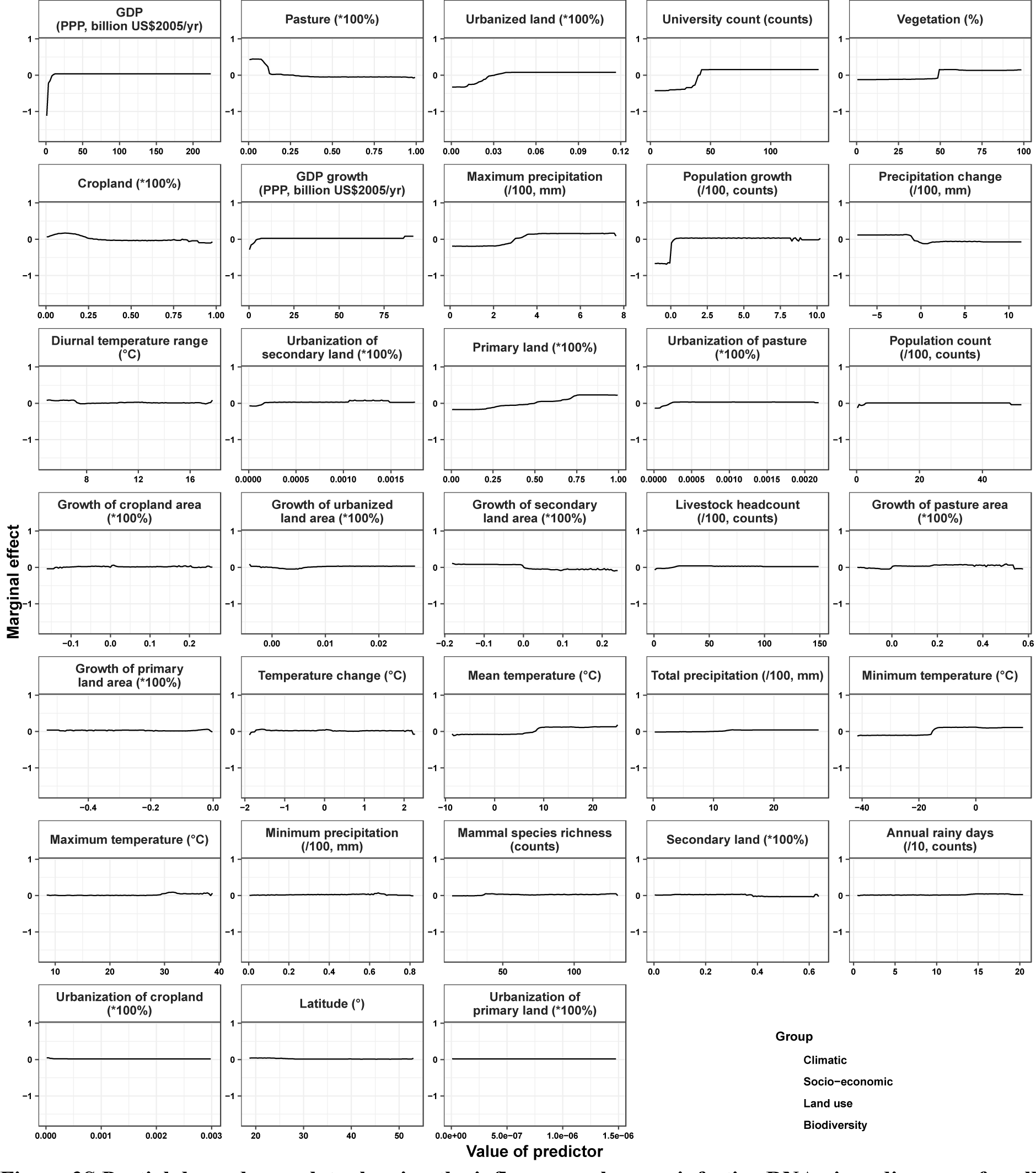
Partial dependence plots showing the influence on human-infective RNA virus discovery for all risk factors in China. Partial dependence plots show the effect of an individual explanatory factor over its range on the response after factoring out other explanatory factors. Fitted lines represent the median (black) and 95% quantiles (coloured) based on 1000 replicated models. Y axes are centred around the mean without scaling. X axes show the range of sampled values of explanatory factors.

**Figure 4S.**
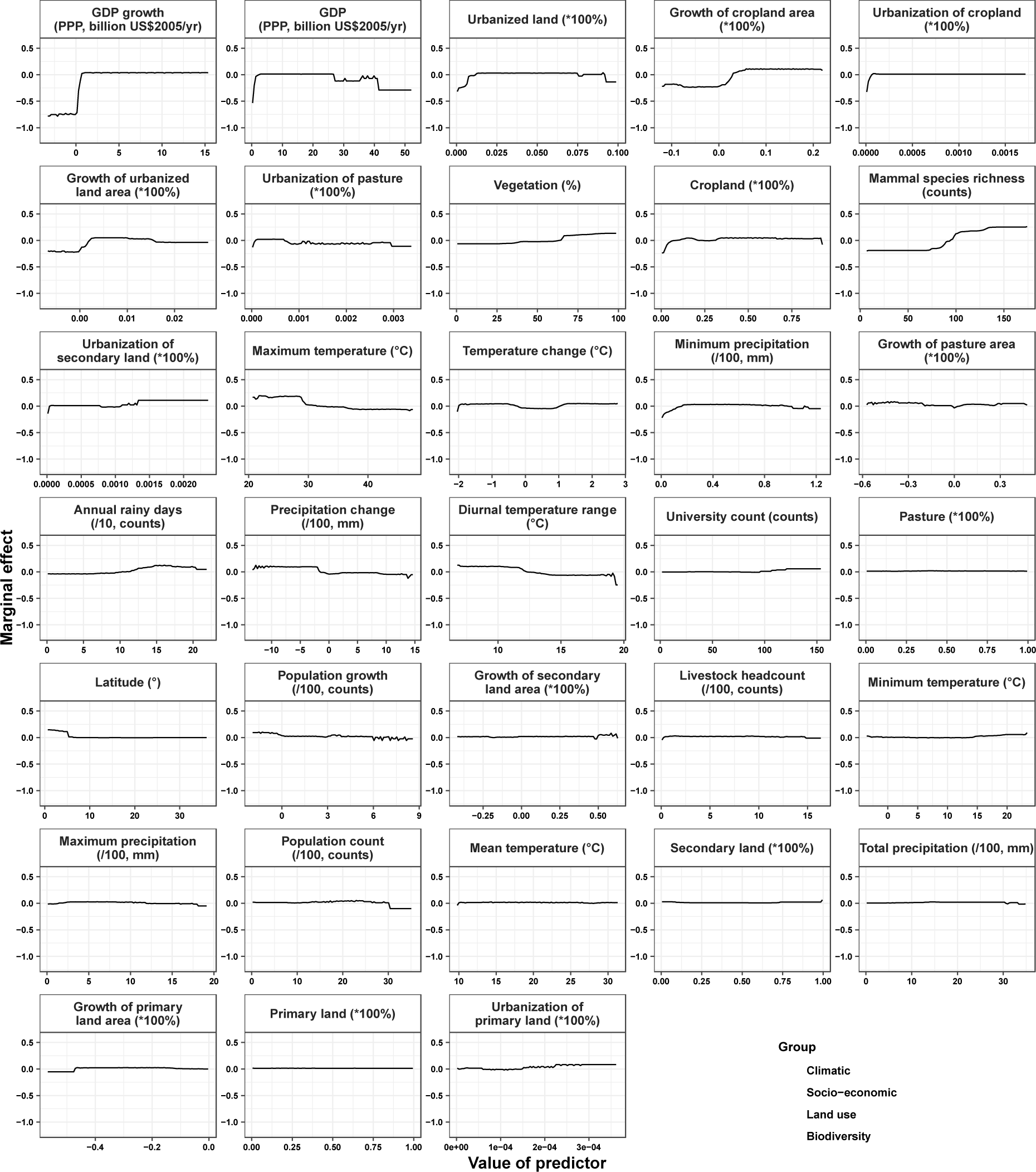
Partial dependence plots showing the influence on human-infective RNA virus discovery for all risk factors in Africa. Partial dependence plots show the effect of an individual explanatory factor over its range on the response after factoring out other explanatory factors. Fitted lines represent the median (black) and 95% quantiles (coloured) based on 1000 replicated models. Y axes are centred around the mean without scaling. X axes show the range of sampled values of explanatory factors.

**Figure 5S.**
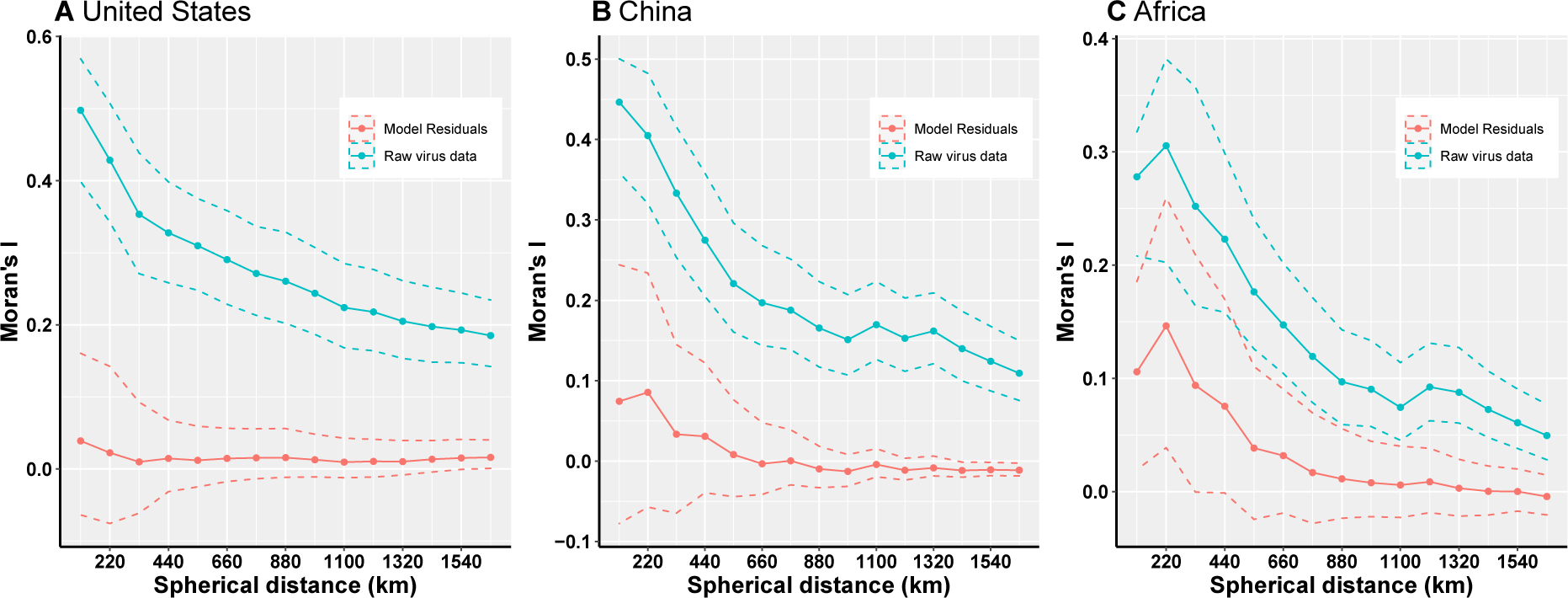
Moran’s I across different spherical distances. A, United States; B, China; C, Africa. The solid line and dots represented the median Moran’s I value, and the grey area represented its 95% quantiles generated from 1000 samples (Blue: Raw virus data) or replicate BRT models (Red: Model residuals). We used the fixed spherical distance as the neighbourhood weights—as there is no general consensus for selecting cut-off values, we chose spherical distances ranging from one time to fifteen times of distance of 1° grid cell at the equator, i.e. 110km to 1650km, considering the area of three regions. Our BRT models reduced Moran’s I value from a range of 0.19–0.50 for the raw virus data to 0.009–0.04 for the model residuals in the United States A), 0.11–0.45 to -0.01–0.09 in China B), 0.05–0.31 to -0.004–0.15 in Africa C), suggesting that BRT models with 33 predictors have adequately accounted for spatial autocorrelations in the raw virus data in all three regions.

**Figure 6S.**
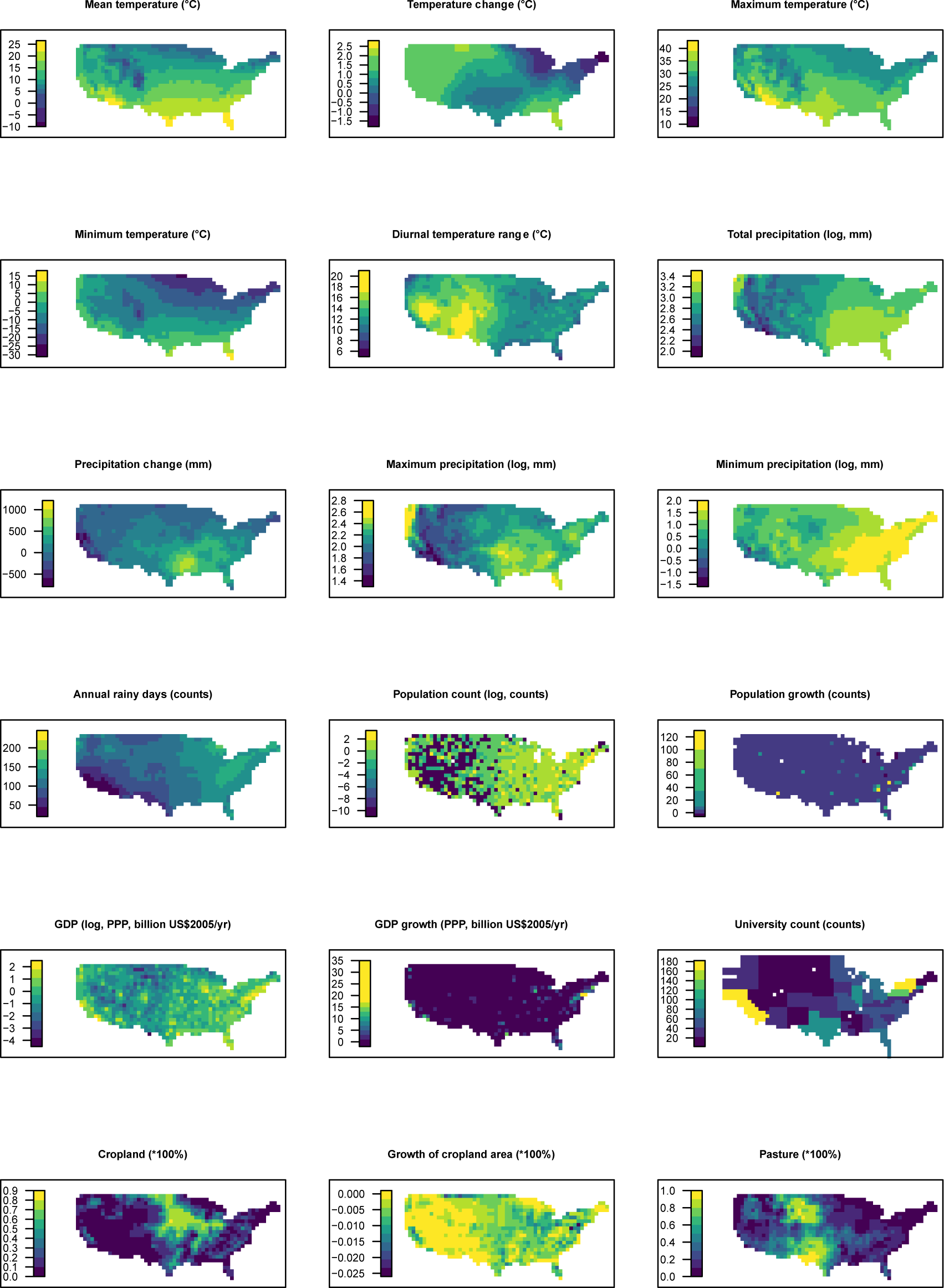
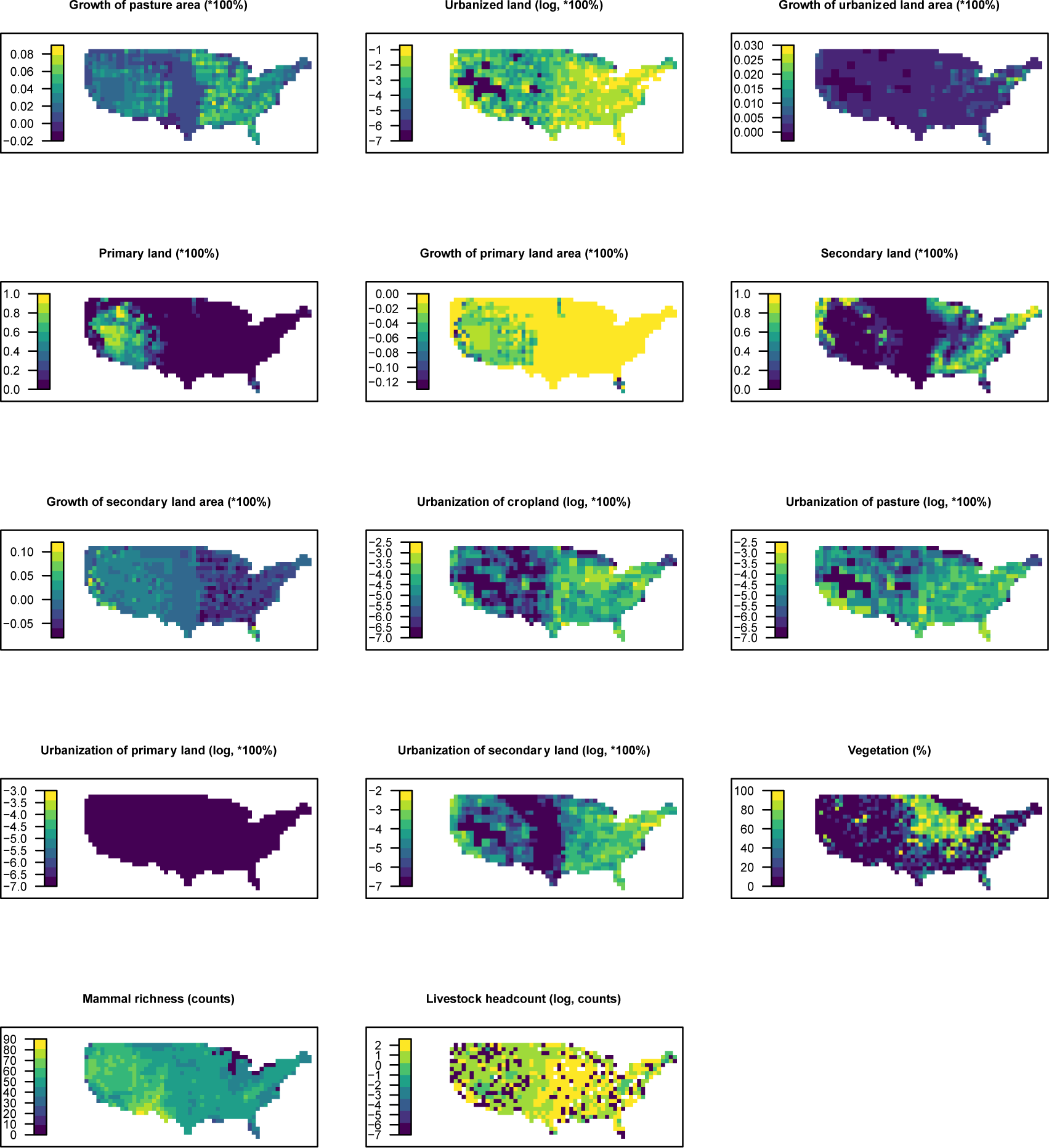
Distribution maps for 32 predictors in 2015 in the United States. The values of these explanatory variables and latitude in each grid cell were used to predict the virus discovery in the corresponding grid cell in the Unites States in 2010–2019. Explanatory variables were log transformed where necessary to get better visualization, not meaning they entered the model by logged values.

**Figure 7S.**
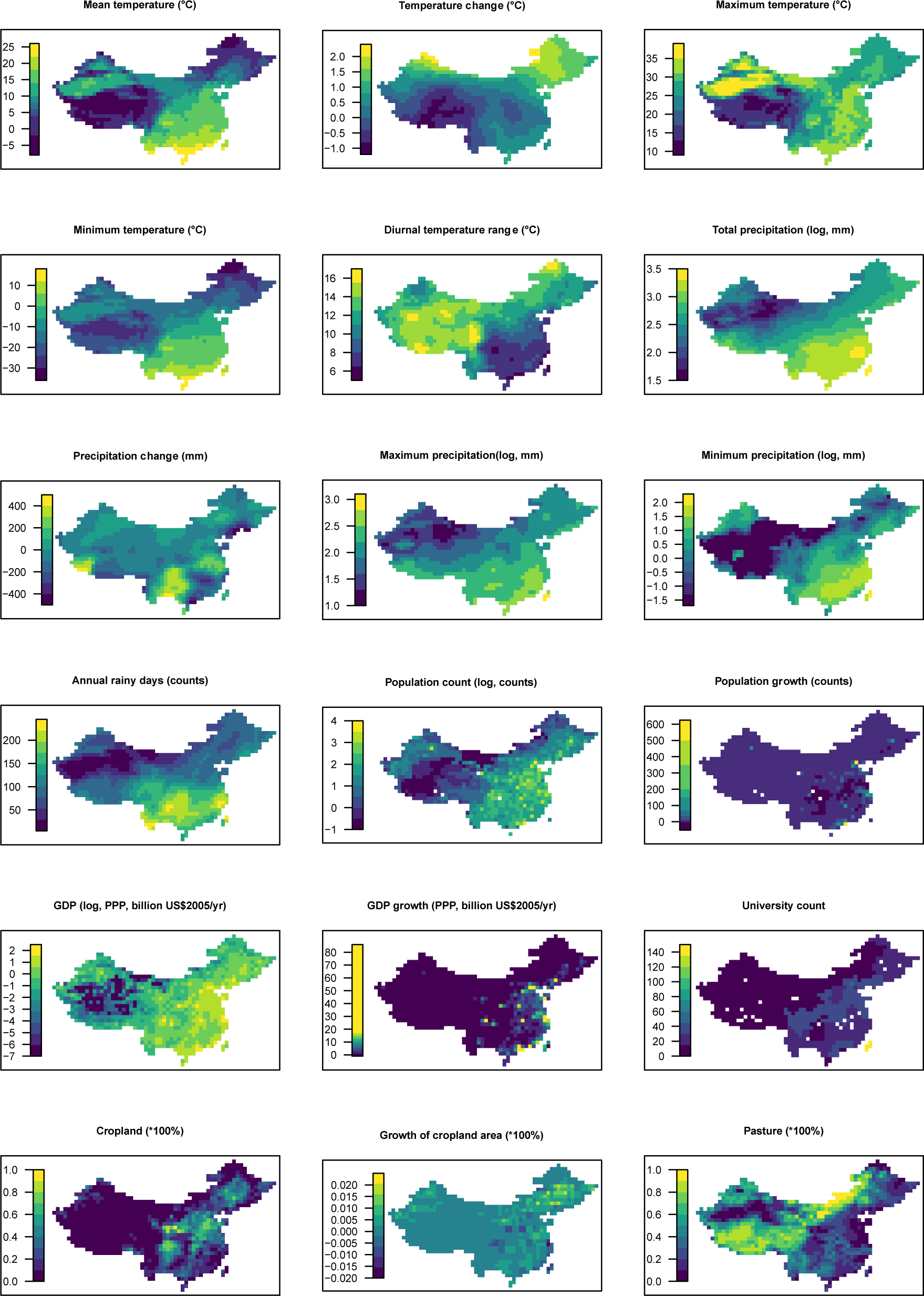
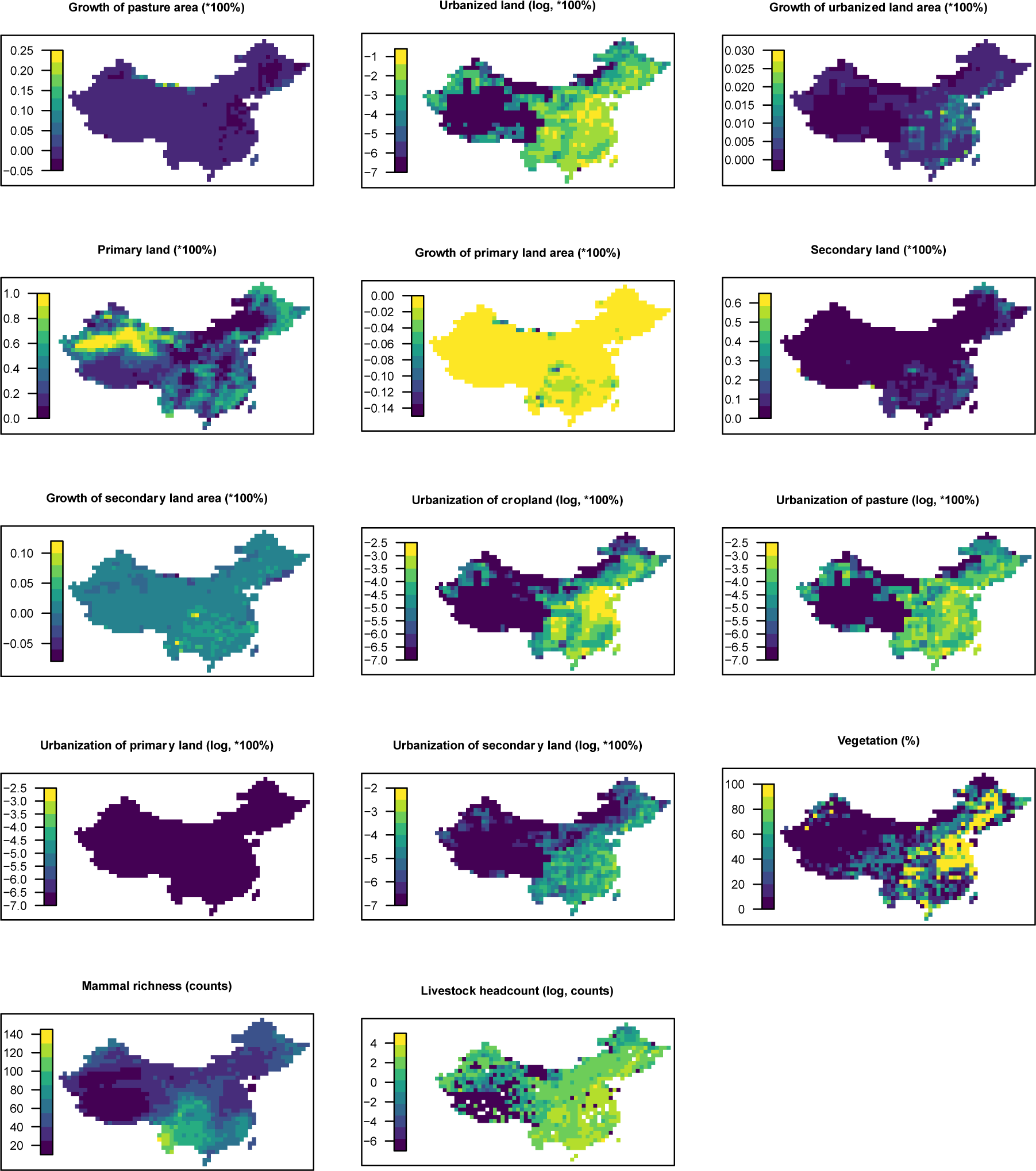
Distribution maps for 32 predictors in 2015 in China. The values of these explanatory variables and latitude in each grid cell were used to predict the virus discovery in the corresponding grid cell in China in 2010–2019. Explanatory variables were log transformed where necessary to get better visualization, not meaning they entered the model by logged values.

**Figure 8S.**
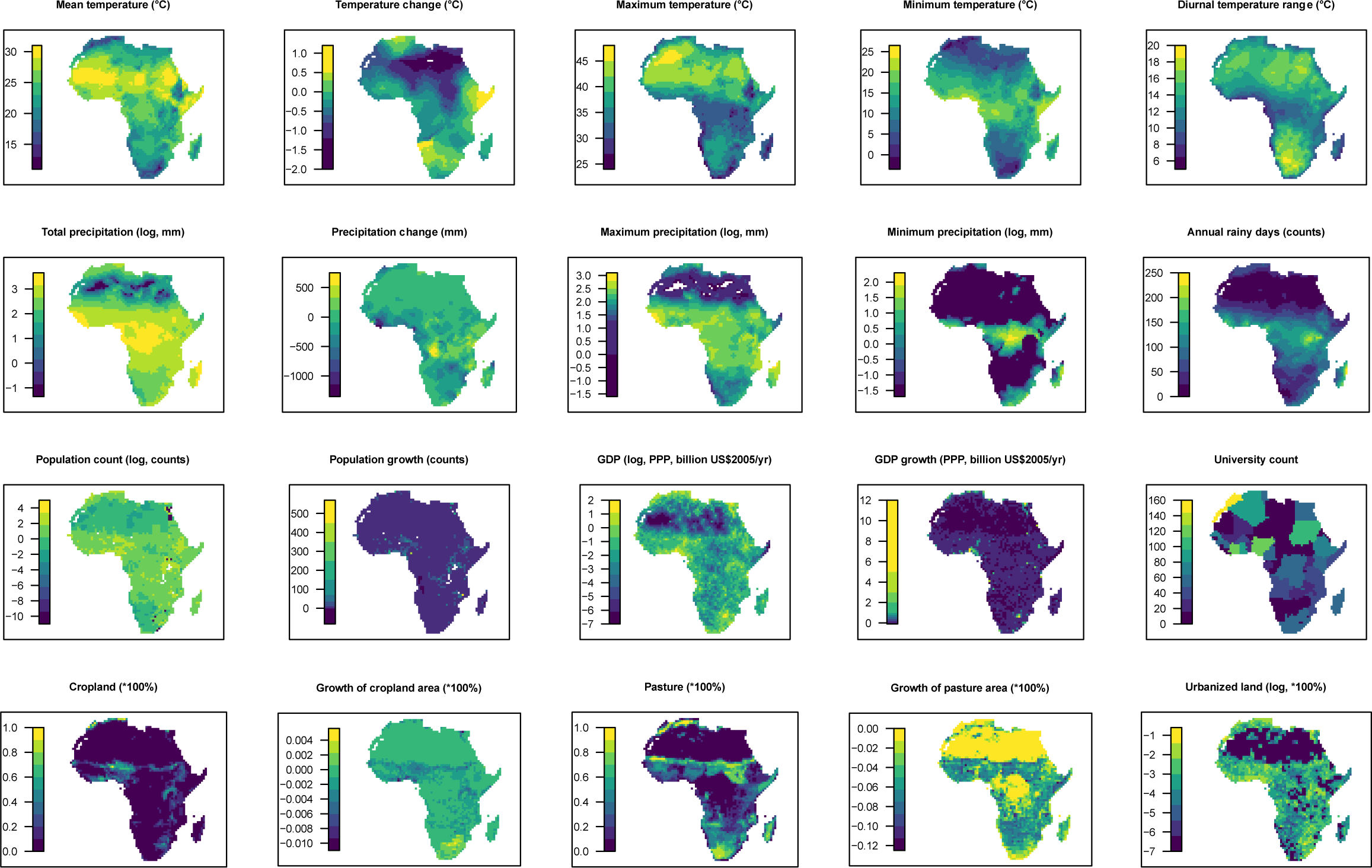
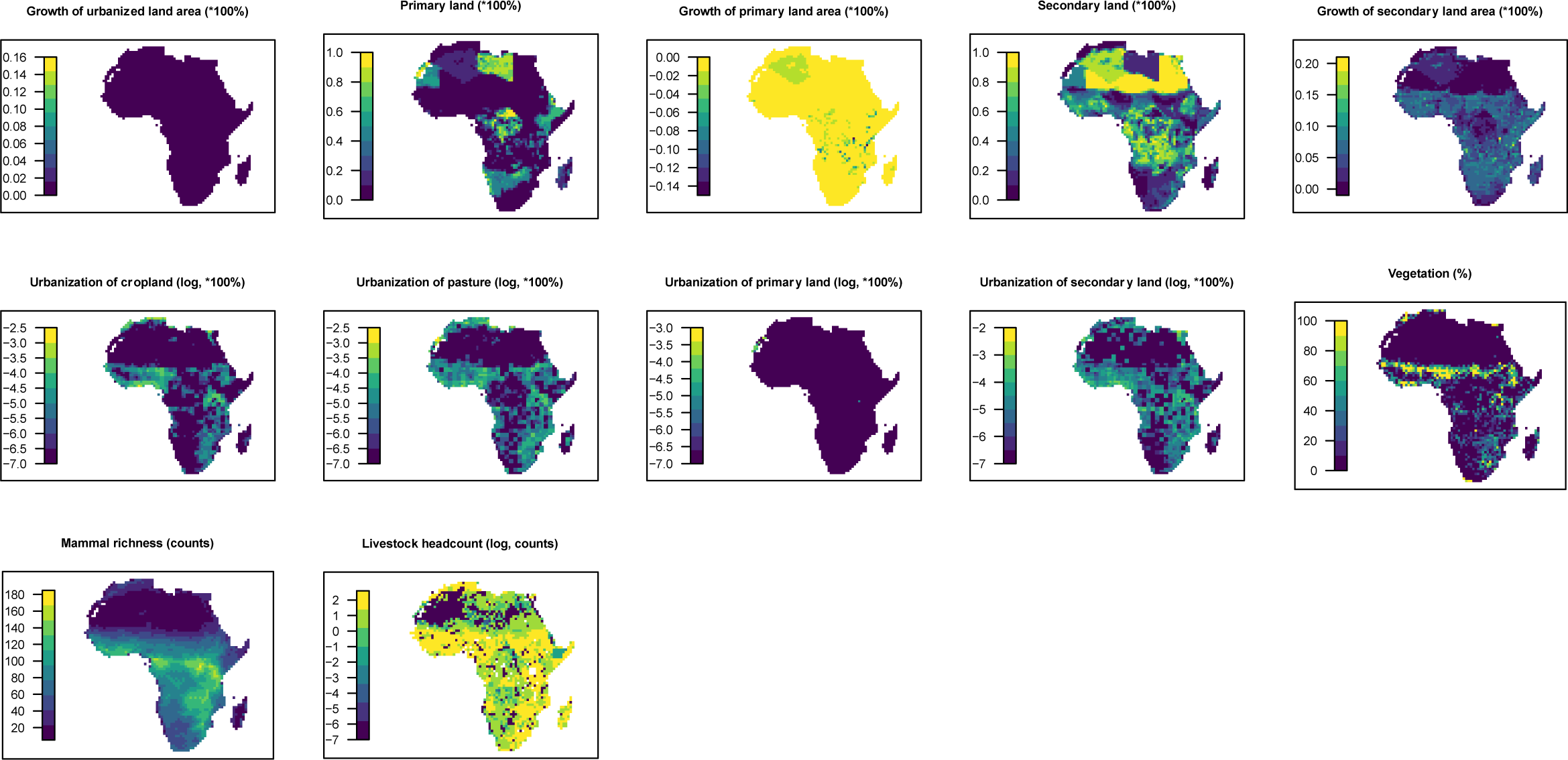
Distribution maps for 32 predictors in 2015 in Africa. The values of these explanatory variables and latitude in each grid cell were used to predict the virus discovery in the corresponding grid cell in Africa in 2010–2019. Explanatory variables were log transformed where necessary to get better visualization, not meaning they entered the model by logged values.

**Figure 9S.**
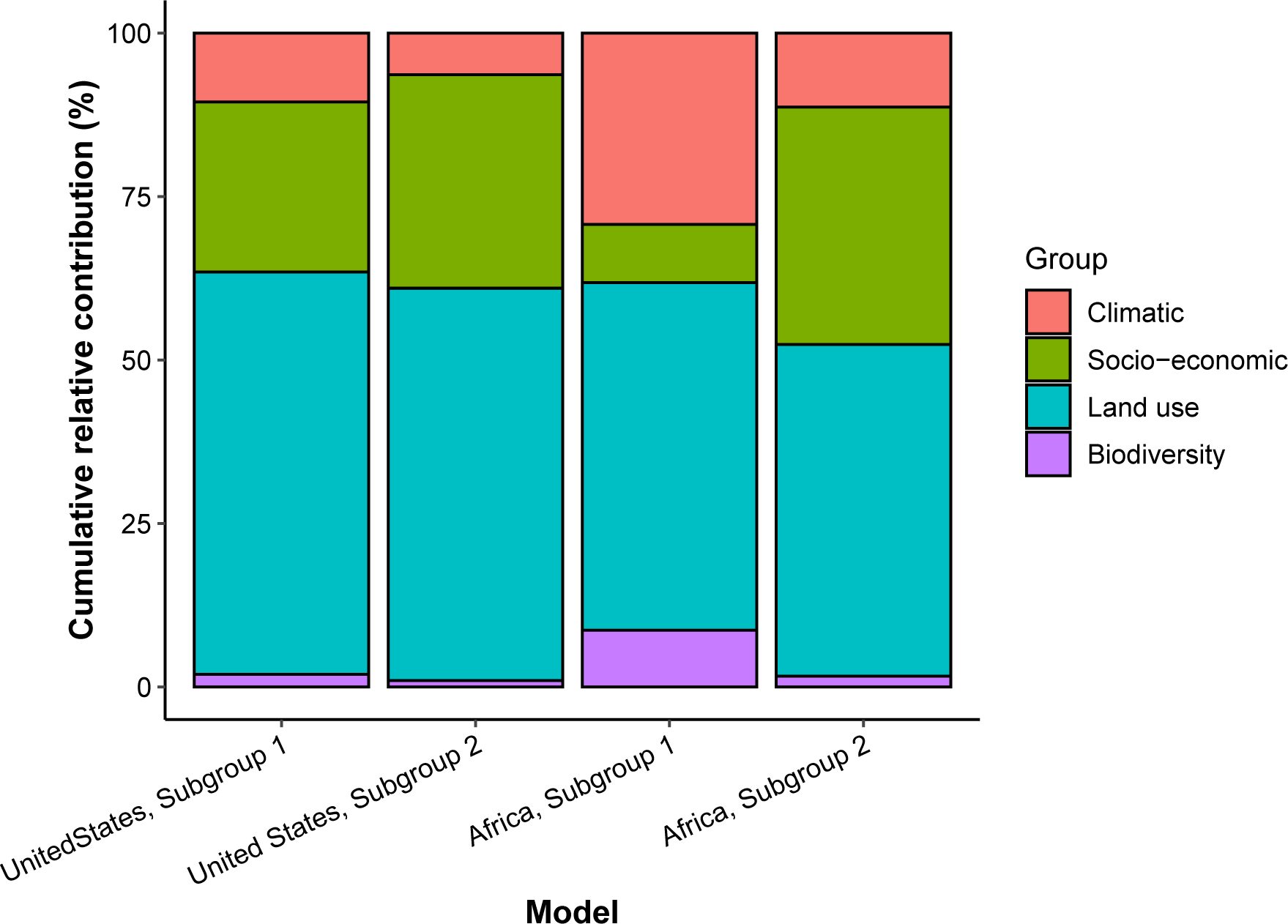
Cumulative relative contribution of predictors to human-infective RNA virus discovery by group in each model of subgroups. Subgroup 1 represents viruses firstly discovered from the region (United States or Africa); Subgroup 2 represents viruses firstly discovered elsewhere in the world. The relative contributions of all explanatory factors sum to 100% in each model, and each colour represents the cumulative relative contribution of all explanatory factors within each group.

